# Revealing the function of HMGB1 N-terminal acetylation by a protein semi-synthesis approach

**DOI:** 10.1101/2021.10.05.463167

**Authors:** Tongyao Wei, Jiamei Liu, Yi Tan, Can Li, Ruohan Wei, Jinzheng Wang, Hongxiang Wu, Qingrong Li, Heng Liu, Yubo Tang, Xuechen Li

## Abstract

HMGB1 (high-mobility group box 1) protein is a nonhistone chromatin-associated protein that has been widely reported to be a representative damage-associated molecular pattern (DAMP) and to play a pivotal role in proinflammatory process once it is in an extracellular location. Accumulating evidence has shown HMGB1 undergoes extensive PTMs that remarkably regulated its conformation, localization, and intermolecular interaction. However, the PTM-related study has been dramatically hindered by the difficulty to access to homogenous proteins with site-specific PTMs of interest. Here, we introduce a protein semi-synthesis strategy via salicylaldehyde ester-mediated chemical ligations (Ser/Thr ligation and Cys/Pen ligation, STL/CPL). This methodology has enabled us to generate N-terminal acetylated HMGB1 proteins in high purity. Further studies revealed that the acetylation on N-terminus regulates its interaction with heparin and modulates its stability, representing a regulatory switch to control the HMGB1’s activity.

## Introduction

High mobility group box 1 (HMGB1) protein is a highly abundant non-histone nuclear protein with critical roles inside as well as outside the cell. Inside the nucleus, HMGB1 as a DNA chaperone and a chromosome guardian can interact with DNA and histones to modify chromatin structures and nuclear processes (e.g., transcription and DNA repair)^1,2,3,4^. Besides, HMGB1 translocation commonly occurs and gains new identities to function in inflammatory or autoimmune disease. Inside the cytoplasm, HMGB1 can be an autophagy sustainer to mediate cell autophagy^5^. Outside the cell, HMGB1 can serve as a key damage-associated molecular pattern (DAMP) and a central mediator of lethal inflammation, facilitating migration and activation of inflammatory cells through Toll-like receptor 4 (TLR4) and advanced glycation end products (RAGE)^1,6^. Also, the extracellular function of HMGB1 can be extended by interacting with pathogen-associated molecular patterns (PAMPs) (e.g., lipopolysaccharide, LPS), cytokines and chemokines^7,8^. By these means, HMGB1 plays multiple roles in the pathogenesis of inflammatory and mediate several important processes ranging from inflammation to repair^5,9^. It is worth mentioning that the relevance of extracellular HMGB1 as a biomarker or therapeutic target has been demonstrated in many diseases^10,11^. For instance, a deleterious systemic inflammatory response (endotoxemia and bacterial sepsis) can be evoked by the circulating bacteria, while it can be subsided by neutralizing extracellular HMGB1, subsequently preventing caspase-11-dependent pyroptosis and death^12^. In the recent COVID-19 pandemic, it was found that serum HMGB1 in severe COVID-19 patents is elevated and plasma concentrations of HMGB1 assessed at ICU admission could accurately identify the fatality for COVID-19 patients^13^. Another study showed that exogenous HMGB1 induces the expression of SARS-CoV-2 entry receptor ACE2 in a RAGE-dependent manner, suggesting a potential target for innovative therapeutic strategies for COVID-19^14,15^.

Over the past few years, the picture of HMGB1 function has grown more nuanced with the recognition that HMGB1 is subjected to extensive posttranslational modifications (PTMs), including methylation, phosphorylation, acetylation, and glycosylation that significantly regulate its activity (e.g., DNA repair and secretion)^4^. A very recent study revealed that OGlcNAcylation at Ser100 in HMGB1 alters its DNA binding on different DNA structures and eventually reduces its DNA damage processing activities^16^. In addition, efficient secretion of HMGB1 requires hyper acetylation on its two nuclear localization sequences (NLS) through JAK/STAT1 pathway, which afterwards causes the accumulation in the cytoplasm^17^. Proteomic studies have identified that in addition to the NLS, significant acetylation was also found at the unstructured N-terminal region (Lys2, 6, 7, and 11) on the extracellular HMGB1 upon cells were exposed to LPS (one of PAMP to stimulate preliminary inflammatory response and HMGB1 release) (Figure S1)^17^. However, to date, a comprehensive investigation of these acetylation in the HMGB1 N-terminus is still lacking.

To study the role of the acetylation at Lys2, 6, 7, and 11, homogenous HMGB1 protein with site-specific lysine acetylation is needed. Towards this end, method modern protein chemical synthesis by combining solid phase peptide synthesis (SPPS) and peptide ligation has evolved to provide versatile strategy to generate proteins with tailor-made posttranslational modifications with precision and flexibility^18,19,20,21,22,23,24^. Here, we reported a highly efficient Ser/Thr ligation mediated semi-synthesis of homogeneous HMGB1 bearing site-specific and different acetylation(s), with which to elucidate the regulatory effect of the acylation on HMGB1’s polysaccharide binding and stability toward enzymatic proteolysis.

A retrosynthetic analysis reveals that HMGB1 protein with the lysine acetylation at the N-terminal region can be assembled via peptide ligation mediated semi-synthesis, in which the short synthetic peptide containing modified residues is ligated with the long peptide fragment without PTMs obtained from recombinant expression^25,26,27,28^. To this end, Thr21-Cys22 could be a potential ligation site for native chemical ligation (NCL), however it might be difficult due to steric hindrance of the beta-branched amino acid^29^. Thus, we explored the HMGB1 semisynthesis via peptide salicylaldehyde (SAL) ester-mediated ligation at the Mer12-Ser13 site. Peptide SAL esters can chemoselectively react with unprotected peptides with N-terminal Ser/Thr (as in Ser/Thr ligation, STL) or Cys/Pen ligation (as in Cys/Pen ligation, CPL) to generate the N,O/S-benzylidene acetal linked intermediate in a pyridine/acetic acid mixture which upon acidolysis restores the natural Xaa-Ser/Thr/Cys/Pen peptide linkage at the ligation site^23,30,31,32,33^. In particular, Cys/Pen ligation is found to proceed independent of the steric hindrance at the C-terminal amino acid (e.g., Val, Ile. Thr and Pro)^33^. Herein, peptide SAL esters-mediated peptide ligation which has not previously been explored for protein semi-synthesis enabled us to efficiently generate HMGB1 proteins with different lysine acetylation at the N-terminal region for the biochemical evaluation.

## Results

### Exemplify STL and CPL in large protein semi-synthesis

Before assembling the HMGB1 protein, we first aimed to establish the working process and optimal conditions to execute peptide SAL ester-mediated peptide semi-synthesis with model peptides/proteins (Figure 1A). As the Ser/Thr ligation uses N-terminal Ser or Thr to mediate the peptide ligation, the first task is to effectively generate the expressed protein with a N-terminal Ser or Thr. The protein with a N-terminal Ser or Thr can be generated through proteolysis by several enzymes, as summarized in Figure S2. Furthermore, proteins with a N-terminal Cys, used for CPL, can also by expressed in a similar manner. Maltose-binding protein (MBP, ∼40 kDa) was chosen for the model study. MBP proteins with N-terminal Ser, Thr, or Cys were obtained by ubiquitin-like-specific protease 1 (Ulp1) digestion. A number of typical peptide SAL esters of different length were prepared via solid phase peptide synthesis (Figure 1B). After extensive condition screening (Table S1), the optimal conditions for STL are identified as following: protein powder was dissolved in hexafluoro-2-propanol (HFIP) or dimethyl sulfoxide (DMSO) at 2 mM and mixed with equal volume of pyridine/acetic acid solution (1:6, v/v); 10 equivalents of peptide ester was added into the solution. Reaction was terminated by ether precipitation after 5 hrs. For CPL, protein powder was dissolved in the phosphate buffer (pH 4) containing 6 M guanidine at 1 mM and 10 equivalents of peptide ester was added into the solution. Reaction was terminated by acetone precipitation after 12 hrs. The STL and CPL intermediates were then subjected to acidolysis with TFA/H_2_O/EDT (90%/5%/5%, v/v/v) for 5-30 minutes and precipitated by ether. The reaction product was analyzed either by SDS-PAGE or LC/MS (Figure 1C, S3, S4, and S5). As shown in Figure 1C, MBP with a N-terminal Thr always had the highest reactivity except for peptides with C-terminal bulky Pro and Val SAL esters. MBP with a N-terminal Cys could react with all the peptide SAL esters smoothly in >60% conversion independent of the C-terminal amino acid. MBP with a N-terminal Ser showed the lowest ligation efficiency but still could finish the reaction in two short peptide esters (P1 and P2) with >95□% conversion based on LC/MS analysis. According to these results, we believed that under the optimized condition, the STL/CPL on large protein semi-synthesis and on short peptides synthesis share the same reaction features (Table S2)^31,33^.

**Figure 1.**
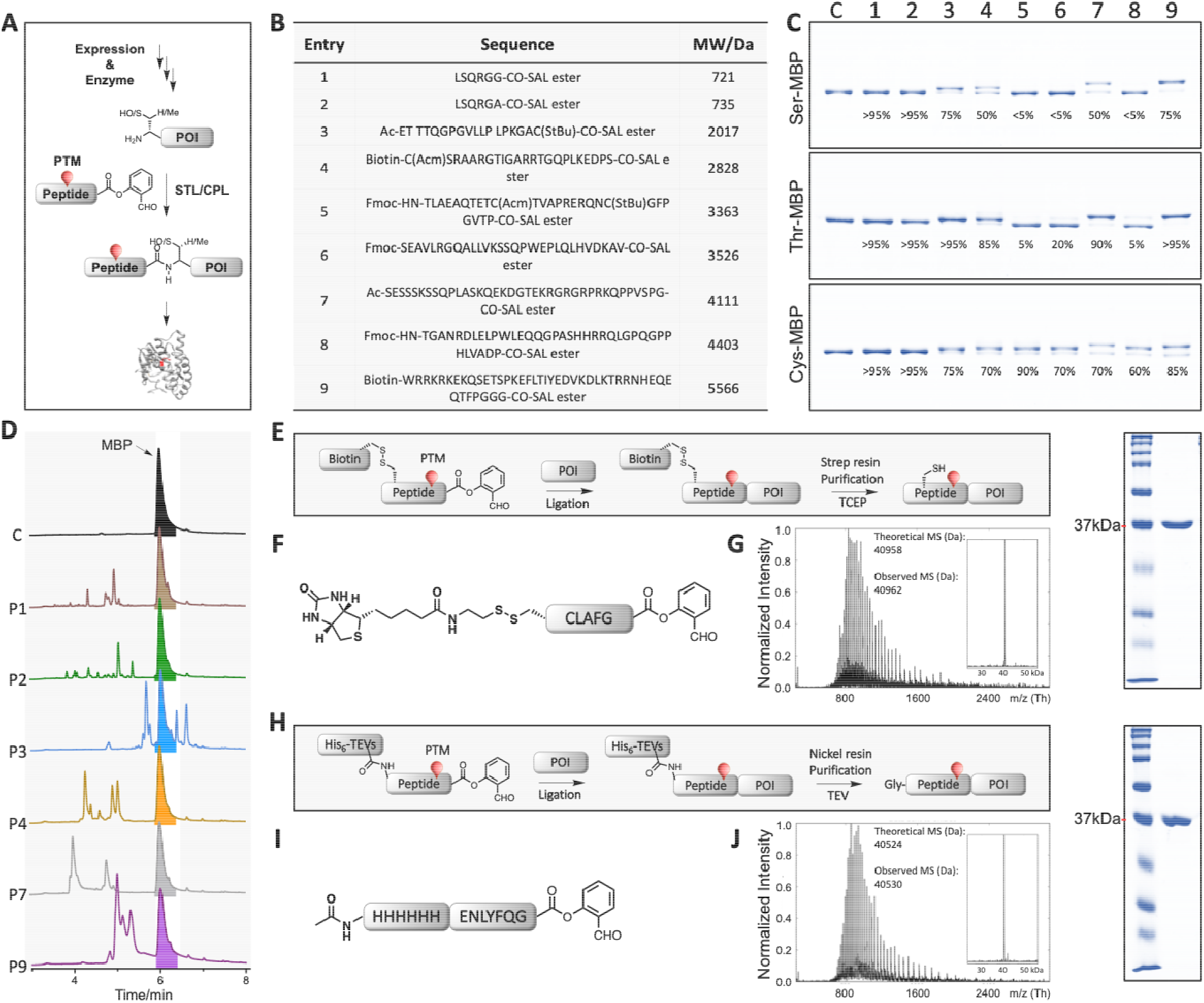
STL/CPL mediated protein semi-synthesis and purification strategies. (A). Overview of STL/CPL mediated protein semi-synthesis. (B). Peptide esters used for demonstrating the ligation. SAL: Salicylaldehyde. (C). SDS-PAGE analysis of ligation products. Conversions were calculated according to the peaks intensity after deconvolution and indicated below the bands. c: control. See also Figure S3, S4, and S5. (D). UV traces of reaction mixtures of the indicated peptide esters reacting with Ser-MBP. The reactions with more than 50% conversion were chosen (Figure 1C top). No shift was observed on the HPLC UV trace after ligation. Except for the product, other peaks mainly were the unconsumed peptide ester, the hydrolysed peptide ester and the product of SAL esters being displaced by HFIP. (E). Disulfide linker-mediated purification strategy. Strep resin: Streptavidin resin. (F). The sequence of model peptide for disulfate linker-mediated purification strategy. (G). Left: Mass spectrum of the final product; Right: SDS-PAGE analysis of the final product. See also Figure S6. (H). His_6_ tag and TEVs based purification strategy. TEVs: TEV protease recognition site. (I). The sequence of model peptide for His_6_ tag and TEVs based purification strategy. Note: This ligation can be fully converted after 3 h. In order to demonstrate our design, we stopped the reaction after 30 min. (J). Left: Mass spectrum of the final product; Right: SDS-PAGE analysis of the final product. See also Figure S6.

As there is a minimal effect on the polarity of the large protein after it is ligated with a short peptide, the final product often overlapped with the unconsumed expressed peptide part on reversed-phase high performance liquid chromatography (HPLC), which made the purification difficult (Figure 1D). To facilitate protein purification, we introduced an extra tag, either His_6_ or biotin tag into the synthetic peptide fragment for subsequent purification. To this end, we used the traceless disulfide linker, which is applicable in cysteine-containing protein. The tag could be removed by tris(2-carboxyethyl)phosphine (TCEP) treatment after purification (Figure 1E, F, and G). Alternatively, the enzymatically cleavable sequence was used, such as tobacco etch virus (TEV) protease recognition site (Figure 1H, I, and J). After purification, the tag could be removed by TEV protease digestion, leaving an extra Gly at N-terminus of the synthetic protein. These two strategies turned out to be very effective to generate homogeneous synthetic proteins, as shown in Figure 1G, and J.

### Semi-synthesis of acetylated HMGB1 proteins

With the protocol established from the above study, we continued with protein semisynthesis of HMGB1 using Ser/Thr ligation strategy (Figure 2A). The synthesis of HMGB1 was designed to disconnect at Met12-Ser13 via Ser ligation. To this end, HMGB1(13-214) was expressed via TEVs-mediated proteolysis (Figure S7), while the peptide (1-12) SAL ester with His_6_-TEV cleavage site and lysine acetylation was chemically synthesized. The ligation proceeded with 35% conversion after 5 hrs according to the deconvoluted peak intensity (the other 65% remain as starting material). After nickel resin purification, TEV protease cleavage, refolding, and size-exclusion purification, the full-length HMGB1 proteins with different lysine acetylation were obtained with 10% overall isolated yield (Figure 2B, and C) and were correctly folded (Figure 2D).

**Figure 2.**
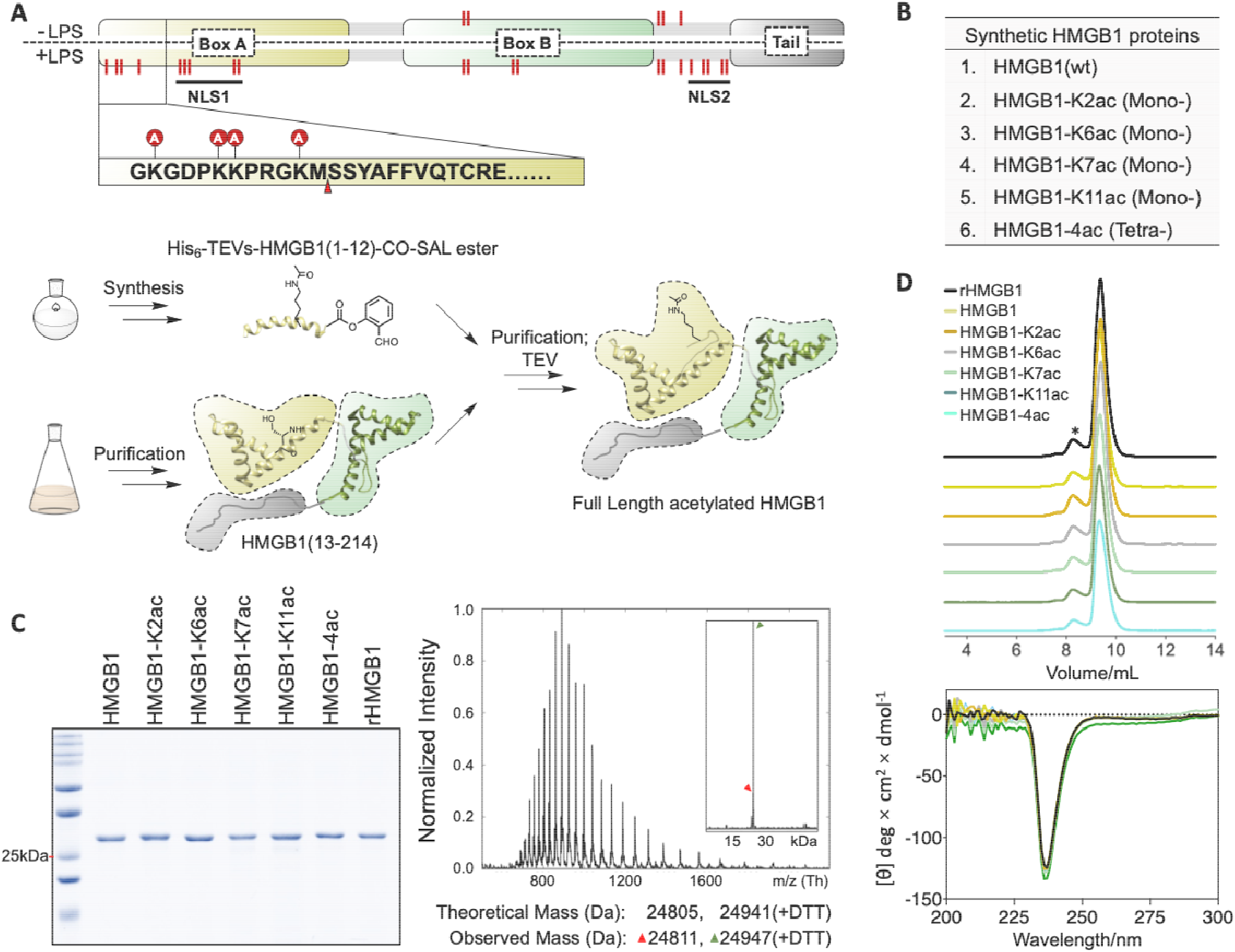
Chemical semi-synthesis of acetylated HMGB1 proteins. (A). Up: HMGB1 was acetylated after LPS stimulation. The modification sites were highlighted by red lines. The ligation site was indicated by an arrowhead. NLS1: first nuclear localization sequence; NLS2: second nuclear localization sequence. Bottom: Semi-synthesis routes of HMGB1. (B). Full length HMGB1 proteins that were synthesized. (C). Left: SDS-PAGE analysis of semi-synthetic HMGB1 proteins and recombinant full length HMGB1 (rHMGB1). Right: Mass spectra of the semi-synthetic HMGB1-K2ac. Note: Intein tag-based purification will install a DTT ester on the C-terminus of protein, see also Figure S7 and S8. (D). Refolding of synthetic HMGB1 proteins. Up: The size-exclusion spectra showed the HMGB1 proteins were folded mainly as monomer; Bottom: Circular dichroism (CD) spectra of HMGB1 proteins. Recombinant full length HMGB1 (rHMGB1) was purified under native condition and was used as a control. *: Dimer of HMGB1.

### Acetylation inhibits HMGB1-heparin interaction

Heparin is an endogenous sulfated glycosoaminoglycan anticoagulant that has been approved for clinical use in the treatment of heart attacks and unstable angina^34,35^. Recently, it was identified as a potent inhibitor of the caspase-11 pathway through targeting HMGB1 and selectively inhibit HMGB1-caspase-11 dependent inflammatory response during sepsis^10,11^. The sequence of HMGB1(6–9) contains a consensus BBXB motif for heparin and other polysaccharides binding, where B is any basic reside and X is any hydrophobic residue^36,37^. To investigate whether the acetylation regulates the HMGB1-polysaccharide interaction, we employed the heparin affinity column, in which the heparin is covalently coupled to highly crosslinked agarose beads. By elution with increasing concentrations of sodium chloride, we determined that all mono-acetylation resulted in decreased heparin-binding, even though the Lys2 and Lys11 do not locate in the BBXB motif, suggesting all four lysines contribute to the surface with positive electrostatic potential of HMGB1 towards heparin binding (Figure 3A). As expected, tetraacetylated HMGB1 was eluted most early due to a synergistic effect (Figure 3A). This results were further validated by sites mutation at these four lysines (Figure 3B) and pull-down experiment using heparin across-linked agarose resin (Figure 3C). In addition, methylene blue (M.B.) was reported as an antagonist of heparin and was used effectively to neutralize heparin and decrease bleeding due to heparinization patients with protamine allergy. Once it binds to heparin, the absorbance peak of M.B. at 664 nm will shift to 568 nm^38^

**Figure 3.**
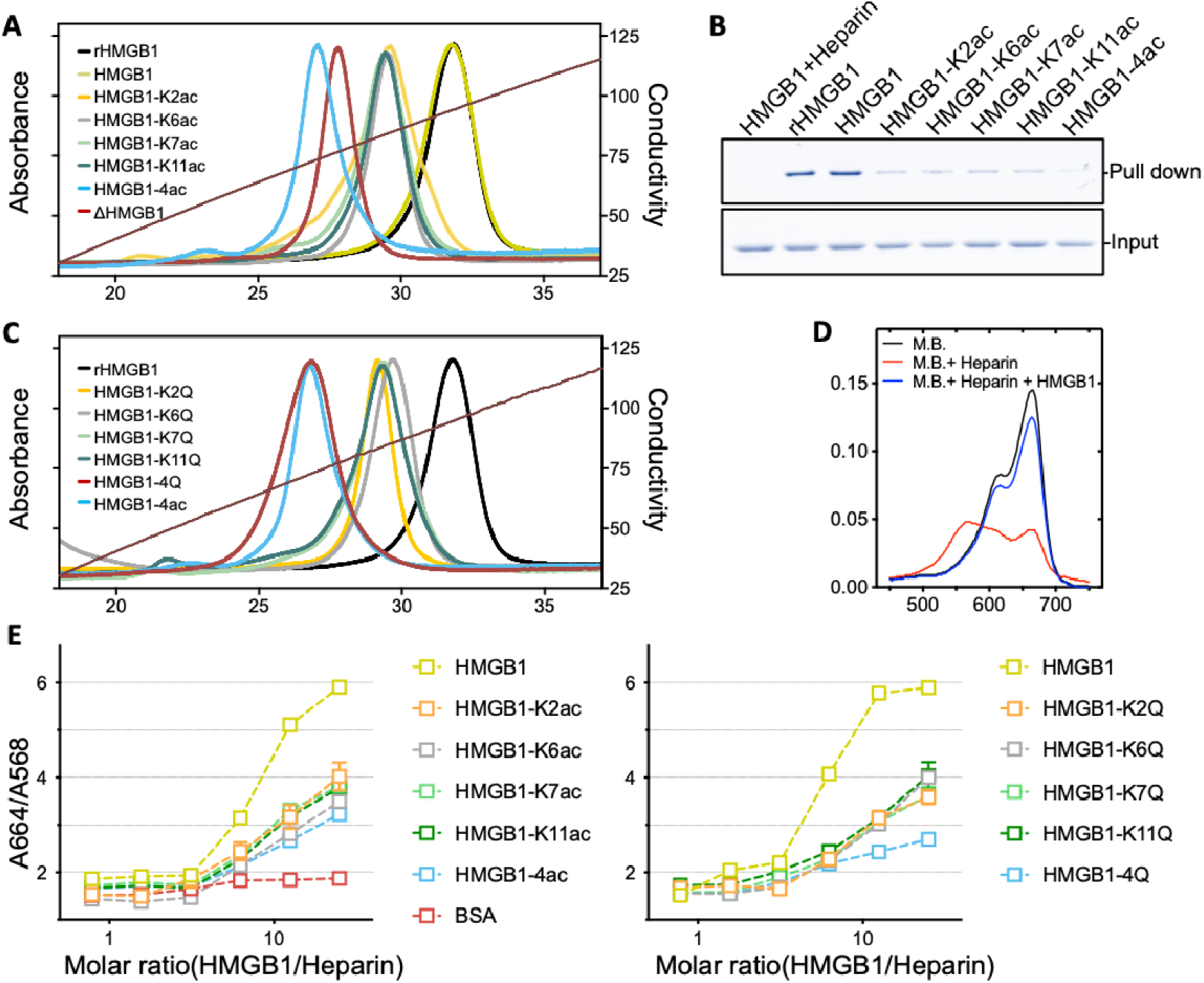
Acetylation is involved in the heparin binding. (A). Heparin column elution spectra of HMGB1 proteins. rHMGB1: full length HMGB1 was purified under native condition. ΔHMGB1: HMGB1 (13-214, N-terminus deleted) was purified under native condition. (B). Pull-down HMGB1 proteins using heparin resin. First lane: Heparin sodium was used as the competitor. (C). Heparin column elution spectra of mutated HMGB1 proteins. (C). Absorbance spectra of M.B., M.B.-heparin complex and M.B. displaced from M.B.-heparin complex by HMGB1. (D). M.B. displacement assay. Left: synthetic HMGB1 proteins; right: mutated HMGB1 proteins. BSA was plotted for negative control. Error bars indicate average ± SEs, n = 3.

However, the addition of HMGB1 restored the absorbance peak at 664 nm, indicating HMGB1 displaced M.B. from heparin effectively (Figure 3D). We also used the ratio between A(664 nm)/A(568 nm) to further evaluate the heparin-binding efficiency of acetylated HMGB1 (Figure S9, Figure 3E left) and mutant HMGB1 (Figure 3E right). Results clearly showed the charge neutralization by acetylation or mutation on N-terminal lysines impairs the HMGB1-heparin interaction.

HMGB1-directed therapies have been tested in several preclinical models, diseases and conditions^10^. The potential benefits of these therapies are well supported in many previous studies^9,10,11,39,40^. In particular, the discovery of sugar based targeting strategies were regarded as a viable avenue to expand the therapeutic window^10^. Going forward, to further defining, improving, and optimizing HMGB1-directed therapies, the intrinsic propensity for posttranslational modification of HMGB1 should be taken into account, as the targeting efficiency can be significantly affected, leading to HMGB1 escape and failure of therapies.

### Acetylation regulates the thrombin-mediated degradation

Thrombin, a serine protease in vascular endothelium, is capable to recognize and cleave the N-terminus at the Arg9-Gly10 site, especially in the presence of thrombin cofactor (thrombomodulin), producing a less proinflammatory form of HMGB1 (Figure S10)^41^. This process has been known as an important mechanism responsible for the rapid clearance of HMGB1 from the circulation^42^. As the N-terminal acetylation sites are adjacent to the cleavage site, we reasoned that these modifications may have a direct influence on the thrombin-mediated degradation. To evaluate the potential roles, an *in vitro* thrombin digestion assay was introduced (Figure 4A). In this assay, a physiological concentration of thrombin (∼1 U/mL), a relatively low concentration of HMGB1 (5 µM), and phosphate-buffered saline were used to reconstitute the extracellular environment. As shown in Figure 4B, thrombin had comparable proteolysis activity on expressed full length HMGB1 (rHMGB1) and synthetic HMGB1 proteins including wild type, Lys2, 7, and 11 acetylation. Remarkably, Lys6 acetylation considerably enhanced the thrombin-mediated cleavage. As a result, tetra-acetylation also facilitated the thrombin digestion.

**Figure 4.**
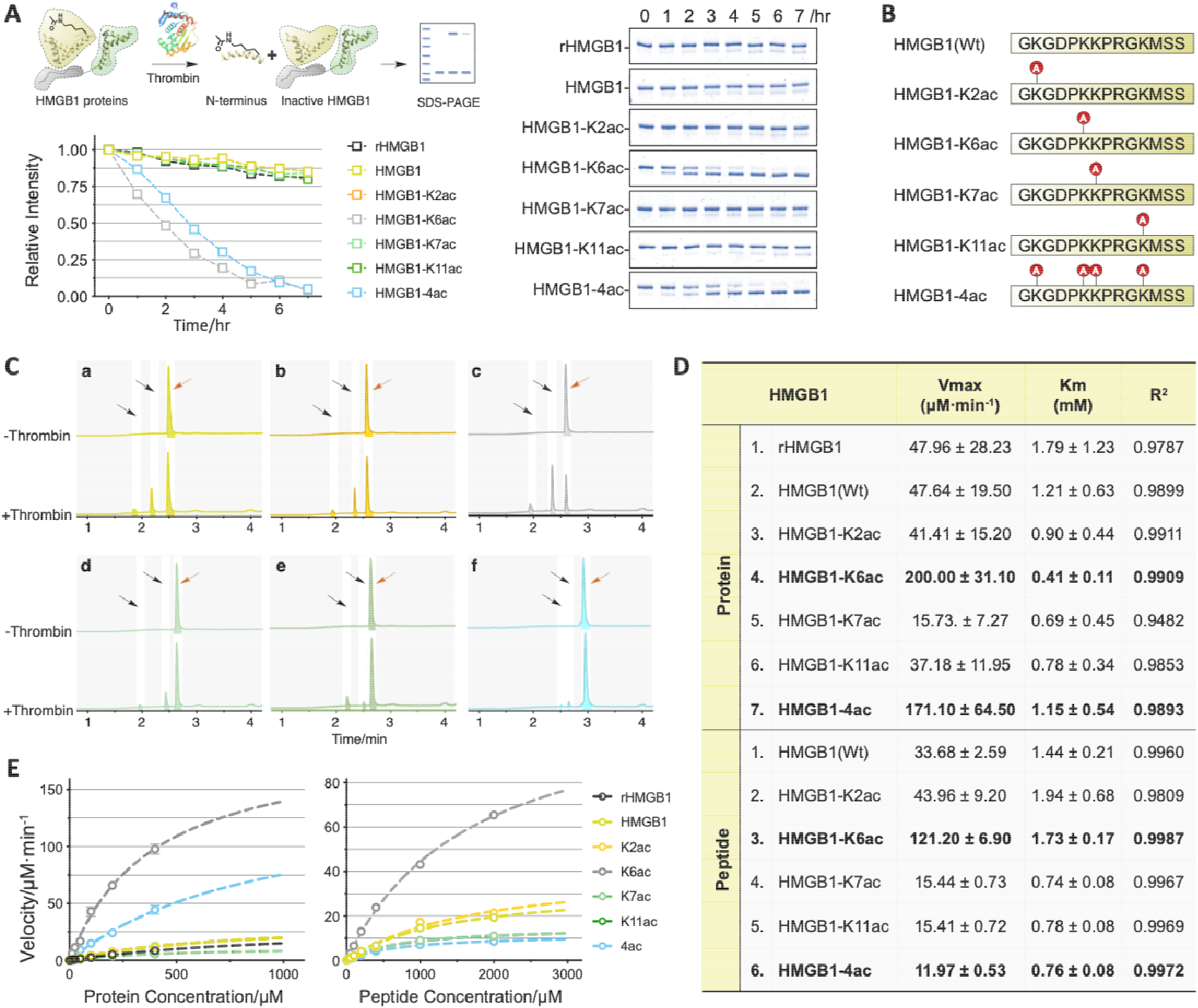
Acetylation is involved in thrombin-mediated degradation. (A). HMGB1 digestion assay. HMGB1 proteins were treated with thrombin and analysed by SDS-PAGE. Digestion rate curve generated from SDS-PAGE (right). Right: SDS-PAGE analysis of digested HMGB1 proteins. (B). Synthetic HMGB1 N-terminal peptides. (C). Thrombin digestion assay on HMGB1 N-terminal peptide. Representative UV traces of the digested HMGB1 N-terminal peptide mixtures after treating with thrombin. a: HMGB1; b: HMGB1-K2ac; c: HMGB1-K6ac; d: HMGB1-K7ac; e: HMGB1-K11ac; f: HMGB1-4ac. (D). Kinetic parameters for thrombin on HMGB1 proteins and N-terminal peptides. Values represent 95% profile likelihood. (E). Michealis-Menten kinetics analysis of thrombin on HMGB1 proteins and N-terminal peptides. Due to protein solubility limits, the highest concentration that we can used is 400 µM.

A previous study has shown that aliphatic residues (*e*.*g*., Leu) at P4 was preferred by thrombin (Figure S11)^43^. As the acetylation on Lys6 (P4) could decrease the hydrophilicity of Lys through installing an acetyl group on its side chain, Lys6 acetylated HMGB1 likely transform into a more optimal substrate for thrombin. To verify our hypothesis, the HMGB1 Nterminus peptides with different acetylation sites were therefore synthesized (Figure 4C). They were then subjected to the thrombin digestion assay. The cleaved products were analysed by HPLC. As expected, since Lys2 is not located in the thrombin, its acetylation barely affects the digestion. Similar to protein, acetylation on Lys6 significantly enhanced the thrombin digestion. In addition, we also observed that Lys7 and Lys11 acetylation slightly hindered the digestion, which was further proved by Michealis-Menten kinetics analysis (Figure D, E, S12, and S13). According to Michealis-Menten kinetics analysis, the acceleration effect from Lys6 acetylation was primarily attributable to an increase in Kcat (Vmax) rather than a decrease in Km, indicating that Lys6 acetylation does not alter the binding affinity towards to thrombin but promote a better position for proteolytic cleavage, while the inhibition from Lys7 and 11 acetylation is caused by a combined effect (both decreased Kcat and Km).

Nevertheless, in addition to Lys7, we also observed that acetylation on Lys11 hindered the digestion, and the overall influence of the tetra-acetylation was negative on peptide digestion, which is opposite to the protein digestion result (Figure 4A, and D). This conflict results prompted us to deduce that the conformational hindrance may play a role to affect the enzyme activity. Indeed, according to HMGB1 boxA’s structure resolved by different groups, the Nterminus especially the region from Pro8 to Met12 is consistently tethered to the third α-helix by several putative hydrogen bonding (Figure S14)^44,45^. Thus, it is very possible that Nterminus is twisted by this intramolecular interaction and is unable to be well positioned for thrombin cleavage like a flexible short peptide. As expected, the side chain of Phe at P4 in a thrombin’s natural substrate (extracellular fragment of human PAR1) has similar direction with the critical Arg at P1 and is in *van der Waals* interaction with Cβ and Cγ of Glu192 of thrombin (Figure S15)^46^. However, we found that the sidechain of Lys11(P4) was stabilized towards to the third α-helix side. Therefore, we inferred that Lys11 sidechain is sequestered by 3D structure and thereby is less possible to actively contribute to thrombin recognition and cleavage, which could explain why the inhibitory effect was not visible on full length HMGB1-K11ac protein. Furthermore, if Lys6 acetylation is capable to counteract Lys7 acetylation’s inhibitory (the overall effect of Lys6 and 7 is still accelerated), tetra-acetylation can lead to an increased digestion eventually.

To validate this, we firstly exam the intramolecular interaction between N-terminus and third α-helix. AlphaFold2 was employed to predict the structures of HMGB1 containing corresponding mutations which are expected to interrupt the putative hydrogen bonds. Recently, AlphaFold2 has been widely used to accurately predict 3D models of protein according to its natural sequence and a high accuracy structure of HMGB1 has been predicted (Figure S14)^47^. As shown in Figure 5A, despite the predicted models are structurally conserved and do not show significant conformational difference between mutations and wild type of HMGB1, we noticed that the prediction confidence for N-terminus was decreased, implying hydrogen bond interruption led to a structural destabilization of N-terminus (low confidence represents a flexible region) (Figure S16)^48^. We also recombinantly generated these HMGB1 mutations accordingly and performed thrombin digestion assay (Figure 5B). An increased digestion was observed in D66A and Y77A mutations, indicating that D66 and Y77 indeed form hydrogen bonds with the N-terminus and contribute to the conformational twist. Abrogating the hydrogen bonds will allow N-terminus to position properly for thrombin cleavage. This conclusion was further proved by HMGB1 truncations digestion. Several HMGB1 truncations were recombinantly generated and subjected to digestion assay (Figure S17). Consistent with sites mutation, we observed significant improvement of thrombin activity when truncating HMGB1 from the third α-helix. Notably, under this theory, thrombin should have higher activity on wild type N-terminus peptide than wild type full length protein, which was not reflected by Michealis-Menten kinetics analysis (Figure 4D). We believed that this error was down to the different quantification methods (SDS-PAGE for proteins, HPLC for peptides).

**Figure 5.**
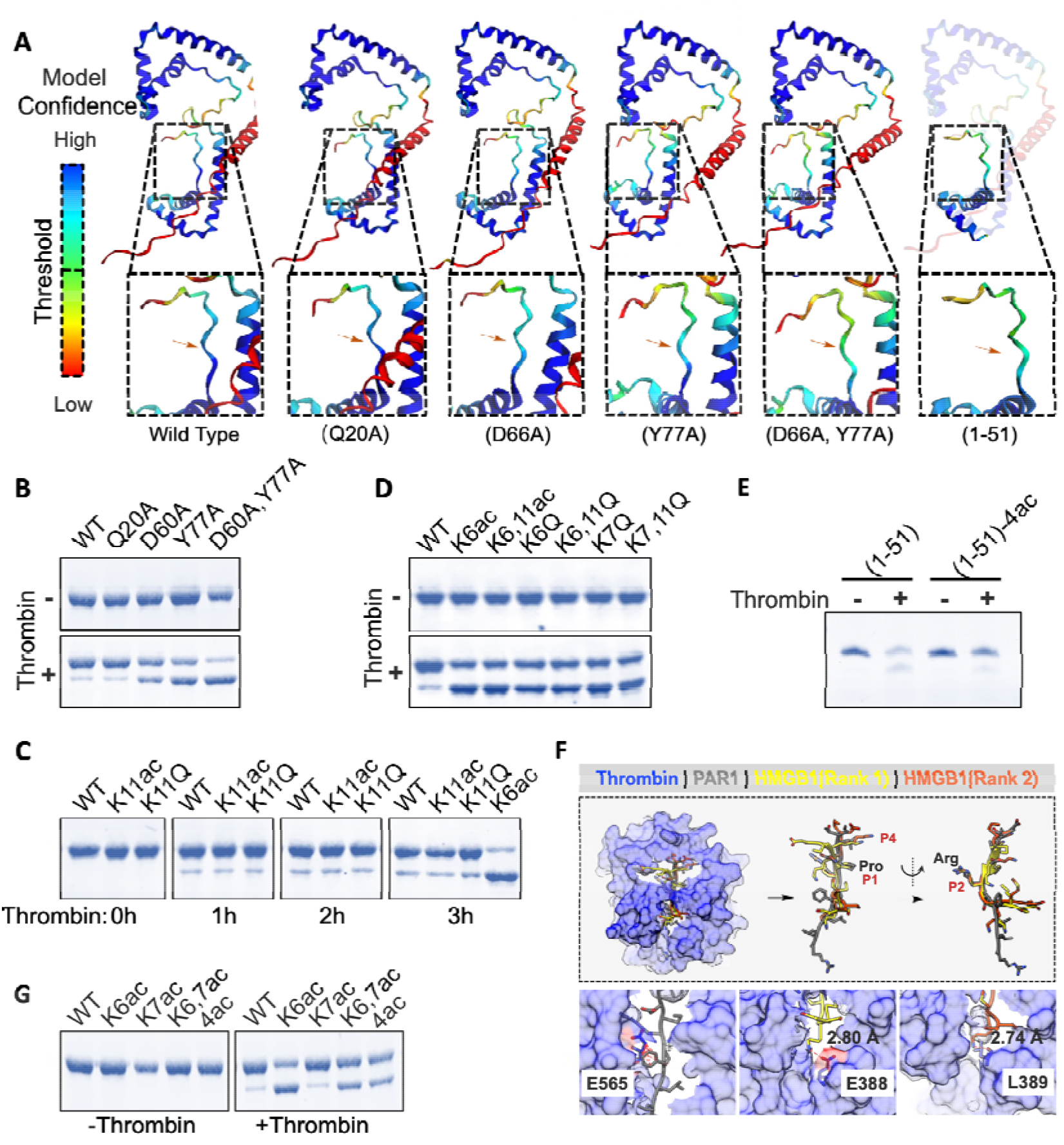
Lys11 is not involved in thrombin-mediated degradation in full length HMGB1 protein. (A). The structures of HMGB1 and its mutations were predicted by AlphaFold2. Structures were coloured by predicted local distance difference test (pLDDT) score (per-residue confidence score). (B). (C). (D). (E). HMGB1 proteins were digested by thrombin. Tricine-SDS-PAGE was used in E. (F). PatchMAN approach for HMGB1 N-terminal docking. Thrombin-PAR1 complex PDB: 3LU9. Thrombin structure from 3LU9 was used for docking. (G). HMGB1 proteins were digested by thrombin.

We next validated that Lys11 sidechain is not involved in thrombin recognition and cleavage on HMGB1 protein. Herein, in addition to HMGB1-K11ac, a Lys11 mutation, HMGB1-K11Q, was introduced, which is regarded as an acetylation mimic. In short peptide, we could observe a significantly inhibitory effect as expected (Figure S18). Importantly, like acetylation, this site mutation still did not show any effect in full-length protein (Figure 5C and S19). Besides, diacetylated HMGB1(K6ac, K11ac) and (K6Q, K11Q) double mutations on HMGB1 were generated. Similarly, the inhibitory effect from K11Q was undetectable (Figure 5D). It is worth mentioning that K7Q mutation, as the mimic of K7ac, yet showed an opposite effect to K7ac, arising the concern about the widely used acetylation mimic. Also, thrombin activity was not affected by on HMGB1 with K7Q, K11Q double mutations (Figure 5D). In addition, we synthesized HMGB1(1-51) with tetra-acetylation (Figure S20). As there is no interference from third α-helix, Lys11 is considered to be functional during thrombin digestion. Unsurprisingly, like N-terminal peptides, tetra-acetylated HMGB1(1-51) was cleaved much slower than its wild type (Figure 5E). We also employed PatchMAN (Patch-Motif AligNments) to simulate the thrombin-HMGB1 N-terminus interaction^49^. While the abilities of computational method for the *de novo* prediction of ligand–protein interactions are limited, there is a main reason why our case is informative and reliable for explaining the interactions. We found that in the top 2 models, HMGB1 N-terminus adopted a conformation similar to that of the native substrate peptide (PAR1) and stably bound to the thrombin in the primary substrate-binding pocket (Figure 5F). In particular, the critical residues (Pro at P2, Arg at P1) in HMGB1 N-terminus match the positioning of that in PAR1 (Figure 5F). The docking results also showed that Lys6, 7, and 11 all contact with thrombin. This could explain why acetylation on these sites affects peptide cleavage. Specifically, the side chains of Lys6 in HMGB1 in both models and Leu38 in PAR1 locate in the same P4 hydrophobic pocket, indicating less hydrophilicity at P4 will benefit thrombin cleavage. Though Lys11 was shown to interact with thrombin in different manner in two models (In rank 1 model, Lys11 interacts with E388 by charge; In rank 2 model, Lys11 forms a backbone hydrogen bond with L389), acetylation or mutation on Lys11 will interrupt both interactions, leading to a decreased thrombin digestion. Above all, we could conclude that in short peptide, Lys11 at P4 is preferred by thrombin while cleavage of HMGB1 protein by thrombin is independent of Lys11 due to the 3D structure.

Finally, we indeed observed that HMGB1 with Lys6 and 7 di-acetylation still showed increased digestion, which indicated Lys6 acetylation is able to outcompete Lys7 acetylation’s inhibitory effect on peptide and protein (Figure 5G, S18). Consequently, the predominant effect from Lys6 acetylation, together with the absence of inhibitory effect from Lys11 acetylation ultimately allow tetra-acetylation to enhance thrombin digestion.

To summarize this section, acetylation on Lys2 does not affect the thrombin digestion; Acetylation on Lys6 greatly improves the thrombin digestion, while acetylation on Lys7 inhibits the thrombin digestion; Lys11 is sequestered by tertiary structure and its acetylation is unable to regulate the thrombin-mediated degradation. Besides, in the cases of multiple acetylation, as Lys2 and Lys7 acetylation do not affect thrombin cleavage, and the inhibitory effect of Lys7 acetylation can be overturned by Lys6 acetylation, Lys6 acetylation always facilitates the digestion independent of the status of other three lysines (*e*.*g*., tetra-acetylation).

In the past decades, extensive signalling crosstalk between phosphorylation and proteolysis have been reported, representing a fine-tuned mechanism to tilt the cell’s fate toward apoptosis^50,51^. However, acetylation, as an another highly abundant posttranslational modification in cells, yet remains largely unexplored in term of this nuanced but important regulation mechanism. Here, the example of Lys6 acetylation-induced thrombin cleavage on HMGB1 will replenish the tool kits of PTMs-directed proteolysis, representing a new multidimensional regulation mechanism in cell activities. On the other hand, the study on Lys11 showed that a short peptide’s structure and function are not necessarily correlated with that of the same sequence in the context of protein, highlighting the importance of protein structural morphism which is capable to sequester PTM-mediated phenotype.

## Discussion

As a sentinel for the immune system, HMGB1 plays a critical role in cell survival/death pathways. In the meaning time, HMGB1 isoforms resulted from posttranslational modifications are also critical for its diverse activities. To systematically elucidate the role of HMGB1 in immune system, a full characterization for the posttranslationally modified isoforms is important. Notably, the main key to explore how the PTMs contribute to the multifarious cellular activities at a molecular level increasingly relies on our ability to access the homogenous proteins with site-specific PTMs of interest. Chemical synthesis offers a useful tool to generate proteins with customized modifications (e.g., PTMs). Thus, a variety of different proteins with complicated modifications (e.g., glycosylation) are becoming accessible^52,53,54^. Nonetheless, the advantage of chemical protein synthesis still remains unfulfilled since the relatively burdensome total synthesis is difficult to start at a biological laboratory. Alternatively, chemical ligation enabled protein semi-synthesis, which combines the strengths of organic synthesis and recombinant protein technology, is more feasible for biologists, as small synthetic peptide parts will be affordable or synthetically accessible by themselves.

In this study, we have established the streamlined peptide SAL ester mediated protein semisynthesis. The success of stitching different peptide SAL esters with the MBP, a 40 kDa-protein, exemplified the operability of STL and CPL on large protein synthesis. Furthermore, two purification strategies were introduced for separating the ligated product from the unreacted or hydrolyzed starting materials without using HPLC purification. In summary, STL/CPL-mediated protein semi-synthesis can serve as a good tool for facilitating the solution to the large protein with PTMs of interest. Using this strategy, we generated multiple full-length HMGB1 proteins with high purity, which enabled us to illustrate that the N-terminal acetylation represents a regulatory switch to control the HMGB1-heparin interaction and HMGB1’s stability. The critical interaction between the HMGB1 N-terminus and polysaccharide (*e*.*g*., heparin) probably has directed the development of sugar-based inhibitors for HMGB1, which is the most popular strategy nowadays^10^. Thus, a detailed study on the N-terminus modification will largely benefit the development of HMGB1-targeted drugs. In addition, thrombin digestion studies revealed a crosstalk between acetylation and circular enzyme-mediated proteolysis, by which acetylation at Lys6 may subverts the proinflammatory function of HMGB1 by accelerating the degradation, and consequently serves as a rescue mechanism from pyroptosis and lethality for cell upon infectious injury (Figure 6).

**Figure 6.**
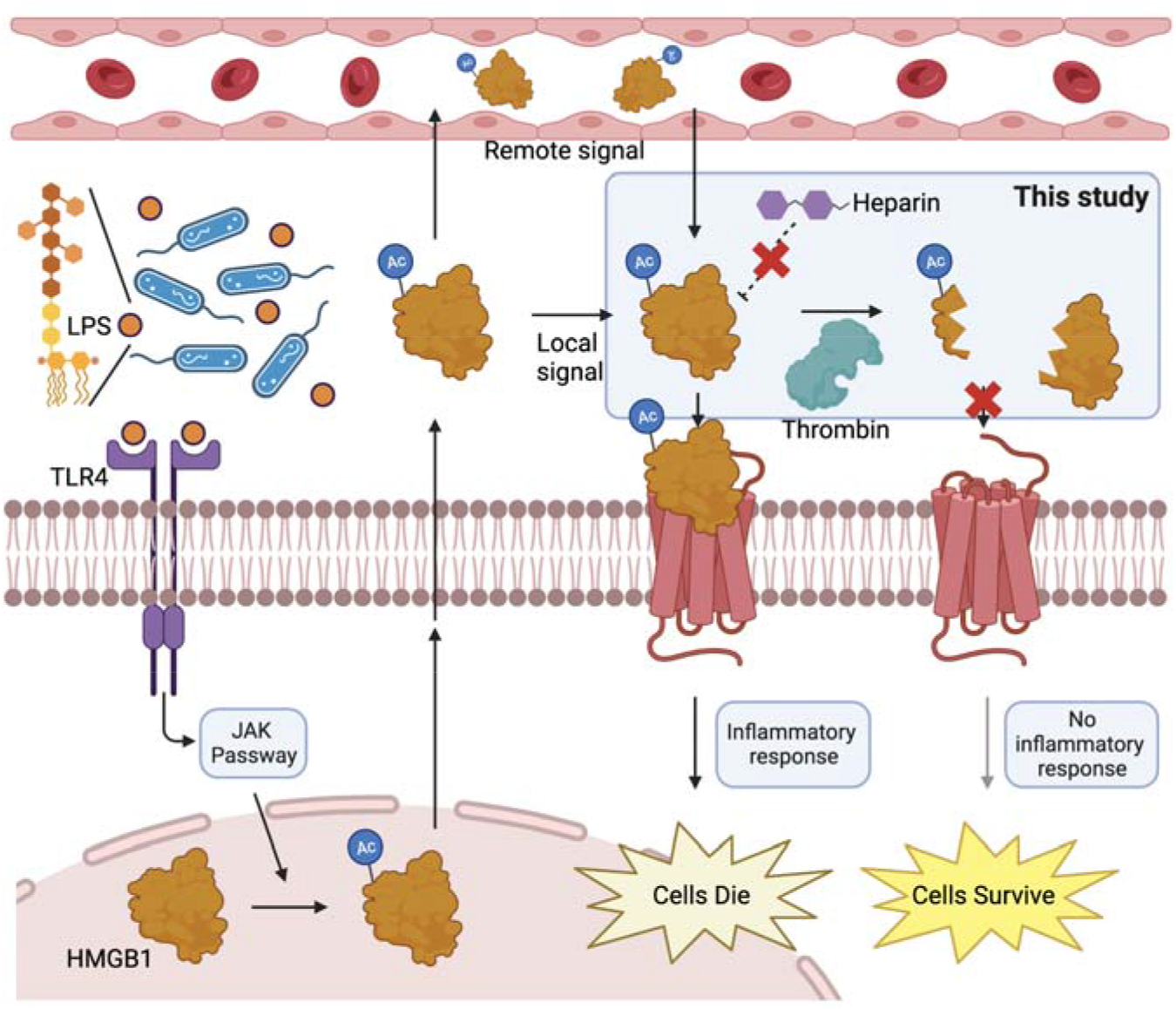
Proposed outcoming for N-terminal acetylation of HMGB1. Circulating bacteria stimulate JAK/STAT1 signalling pathway. Afterwards, HMGB1 is hyper acetylated including the N-terminal acetylation and is secreted. In extracellular, N-terminal acetylation impair the targeting efficiency of heparin. On the other hand, the acetylation at Lys6 promotes HMGB1 to be degraded rapidly by thrombin. As a result, acetylated HMGB1 fails to stimulate more severe inflammatory responses, and cells survive.

Please note that this manuscript has been preprinted at bioRxiv: (https://www.biorxiv.org/content/10.1101/2021.10.05.463167v1.full) It has been cited by (*Chem. Sci*., 2022, **13**, 1367-1374), in which a new HMGB1 protein semisynthesis approach was developed based on figure 1F of this study and a solubilizing tag.

## Methods

### Plasmids

MBP DNA was cloned into pET28a with an N-terminal SUMO tag to generate pET28a-His-SUMO-Ser-MBP, pET28a-His-SUMO-Thr-MBP, and pET28a-His-SUMO-Cys-MBP plasmids for Ser-MBP, Thr-MBP and Cys-MBP expression, respectively. HMGB1 sequence was optimized for *E. coli* codon and the HMGB1 mutations and truncations DNA was inserted into PET28a with an N-terminal TEV recognition sequence and C-terminal inteinCBD-His_6_ tag to generate pET28a-TEV-HMGB1s-intein-CBD-His plasmid for HMGB1 expression.

### Protein expression and purification

All pET28a plasmids were transformed into BL21. The expression of MBP proteins was induced by including 0.2 mM isopropyl β-D-1thiogalactopyranoside (IPTG) when OD600 reached 0.8, and the culture was grown at 25 °Cfor overnight. Bacteria were collected by centrifugation then sonicated in lysis buffer (50 mM Tris-HCl, pH 7.5, 500 mM NaCl, 50 mM imidazole, 1mM PMSF). After centrifugation, the supernatant was loaded onto the Histrap HP column, followed by thoroughly washing with 50 mL lysis buffer. The MBP proteins were eluted by 10 mL elution buffer (50 mM Tris-HCl, pH 7.5, 500 mM NaCl, 500 mM imidazole). After digestion by Ulp1 to release N-teminal Ser, Thr and Cys, the solution was directly injected into HPLC for Ser/Thr/Cys-MBP purification (10-40% CH_3_CN/H_2_O over 40 min). Finally, the eluted fractions were checked by LC-MS, and the desired fractions were combined and lyophilized to afford the protein powder. For HMGB1 purification, initially, the His-TEV-HMGB1 constructions produced several truncated species probably due to the highly negative charged acidic tail (Figure S7). Therefore, the His_6_ tag was inserted at the C-terminus. In addition, to minimize the extra sequence on HMGB1 protein after purification, the intein-CBD tag was inserted between HMGB1 and the His_6_ tag. The expression was induced by including 0.2 mM IPTG when OD_600_ reached 0.8, and the culture was grown at 16 °Cfor overnight. The bacteria were collected by centrifugation then sonicated in lysis buffer (50 mM Tris-HCl, pH 7.5, 500 mM NaCl, 50 mM imidazole, 1mM PMSF). After centrifugation, the supernatant was loaded onto the Histrap HP column, followed by thoroughly washing with 50 mL lysis buffer. The HMGB1 protein was eluted by 10 mL elution buffer (50 mM Tris-HCl, pH 7.5, 500 mM NaCl, 500 mM imidazole). After digestion by TEV to release the N-terminal Ser, 100 mM DTT was included to trigger intein splicing. After 24 h, the solution was directly injected into HPLC for HMGB1(13-214) purification (10-40% CH_3_CN/H_2_O over 40 min). The eluted fractions were checked by LC-MS. The fractions containing HMGB1-DTT and the hydrolysis product of HMGB1-DTT were combined and lyophilized. The full length HMGB1 and truncations were purified following the same procedure.

### Peptide and peptide salicylaldehyde (SAL) esters synthesis

Peptides were synthesized under standard Fmoc/tBu SPPS protocols. One equivalent of trityl resin was reacted with four equivalents of amino acids and coupling reagent in DMF. HATU/DIPEA was used for all the peptide coupling steps. The deprotection solution was the mixture of piperidine/DMF (1/4, v/v). The details can be found in supplementary information.

### Chemical ligation

All ligations were set up at room temperature. HFIP or DMSO was used as the cosolvent to dissolve proteins firstly. For STL, Ser/Thr-MBP powders were dissolved in HFIP at 2 mM, then mixed with equal volume of pyridine/acetic acid solution (1:6, v/v); 10 equivalents of peptide esters were added into the solution. After 5 h, reactions were terminated by adding 10-fold (volume) of ether to precipitate proteins and peptides. For CPL, Cys-MBP powder was dissolved in 6 M Guanidine in phosphate buffer (pH = 4) at 1 mM and 10 equivalents of peptide esters were added into the solution. After 24 h, the reaction was terminated by adding 4-fold (volume) of acetone to precipitate proteins and peptides. The precipitates were collected by centrifugation. After being dried by the stream of flow air, the STL and CPL intermediates were then subjected to acidolysis with TFA/H_2_O/EDT (90%/5%/5%; v/v/v) for 5-30 minutes and precipitated by ether again. The final products were analyzed by LC/MS and SDS-PAGE. The HMGB1 (13-214) powder was dissolved in DMSO at 2 mM, then mixed with equal volume of pyridine/acetic acid solution (1:6, v/v); 5 equivalents of His_6_-TEV-HMGB1(1-12) esters were added into the solution. After 5 h, the reactions were terminated by adding 10fold (volume) of ether to precipitate proteins and peptides. The precipitates were collected by centrifugation. After being dried by the stream of flow air, the STL and CPL intermediates were then subjected to acidolysis with TFA/H_2_O/EDT (90%/5%/5%, v/v/v) for 15 minutes and precipitated by ether again. The solids were collected. Of note, the HMGB1 related salicylaldehyde esters were unexpectedly found to undergo hydrolysis quickly in HFIP, resulting in poor yield (less than 10%). Thus, DMSO was used as cosolvent instead. However, the ligation efficiency in DMSO was slightly lower than in HFIP for other peptide SAL esters. In summary, for selection of cosolvent, DMSO was usable for all tested peptide SAL esters. HFIP, which could give higher conversion in most cases, while may cause hydrolysis of peptide SAL esters in some cases.

### Disulfide linker mediated purification

After chemical ligation, the solids were dissolved in 6 M guanidine, followed by 10 times dilution in refolding buffer (50 mM Tris-HCl, pH 7.5, 500 mM NaCl). After centrifugation, the supernatant was loaded onto the strep resin. After complete washing, the protein was eluted by directly treating resin with 20 mM TCEP for 1 h. The remining peptides can be removed by following concentration step.

### His_6_ tag and TEV based purification

After chemical ligation, the solid was dissolved in 6 M guanidine, followed by 10 times dilution in refolding buffer (50 mM Tris-HCl, pH 7.5, 500 mM NaCl). After centrifugation, the supernatant was loaded onto nickel resin. After complete washing, the protein was eluted by elution buffer (50 mM Tris-HCl, pH 7.5, 500 mM NaCl, 500 mM imidazole). The eluted fraction was desalted and digested by TEV. After digestion, the protein solution was re-loaded onto the nickel resin. The flowthrough was collected. The remining peptides could be removed by following concentration step.

### HMGB1 refolding, re-purification

After chemical ligation, the solid was dissolved in 6 M guanidine, followed by 10 times dilution in refolding buffer (50 mM Tris-HCl, pH 7.5, 500 mM NaCl). After centrifugation, the supernatant was loaded onto size-exclusion column. The according HMGB1 monomer fractions were collected and further purified by nickel resin. The eluted fraction was digested by TEV to give the full length HMGB1proteins.

### Heparin column elution

HMGB1 proteins were exchange in 10 mM sodium phosphate (pH ∼7, no NaCl) and loaded on HiTrap™ Heparin HP (5 mL) column. After washing with 5 mL sodium phosphate (pH ∼7), HMGB1 proteins were eluted with 10 mM sodium phosphate (pH ∼7) containing 1M NaCl using a continuous gradient from 0%-100% in 25 mL.

### Pull-down experiment

20 µg HMGB1 proteins was incubated with 20 µL heparin resin slurry in binding buffer (20 mM Tris-HCl, pH 7.5, 100 mM NaCl, 10% glycerol, 0.2 mM EDTA, 0.1% Triton X-100) for 1 h. After twice washing by binding buffer, the resin was resuspended in 40 µL 1x loading buffer for SDS-PAGE resolving and Coomassie blue staining.

### Thrombin digestion

HMGB1 proteins were diluted in PBS at 5 µM, then treated with thrombin (1 U/mL) at 37 °C. The digestion was monitored at indicated time points (Figure 4A and Figure 5). 20 U/mL thrombin was used for Michealis-Menten kinetics analysis. HMGB1 N-terminal peptides were diluted in PBS, then treated with thrombin (20 U/mL) at 37 °C. The digestion was monitored by HPLC at indicated time points.

### Computational predication

The structures of HMGB1 and its mutations/truncations were predicted by AlphaFold2 online server: ColabFold (https://colab.research.google.com/github/sokrypton/ColabFold/blob/main/AlphaFold2.ipynb); Thrombin and HMGB1 N-terminus complex structures were predicted by PatchMAN online server (https://furmanlab.cs.huji.ac.il/patchman/job/f3ac2df5-5f06-4d9c-be0c-6ded640583e9).

## Acknowledgements

The work was supported by the Grants Research Council of University Grants Committee of Hong Kong (C7147-20G, 17303920, AoE/P-706/16). We thank Dr. Han Liu for proofreading the manuscript.

## Author Contributions

X.L., T.W. designed the experiments, analyzed the data, and wrote the paper with input from other authors. T.W. and C.L. performed biochemistry experiments. T.W., J.L., Y.T., J.W., and H.W. performed synthetic experiments. R.W., Q. L., H.L. and Y.T. provided discussion.

## Additional information

**Supplementary information** The online version contains supplementary material available at **Correspondence** and requests for materials should be addressed to Prof. Xuechen Li.

