## Supplementary material for "Revealing the function of HMGB1 N-terminal acetylation by a protein semi-synthesis approach": SI

### Supplementary Information

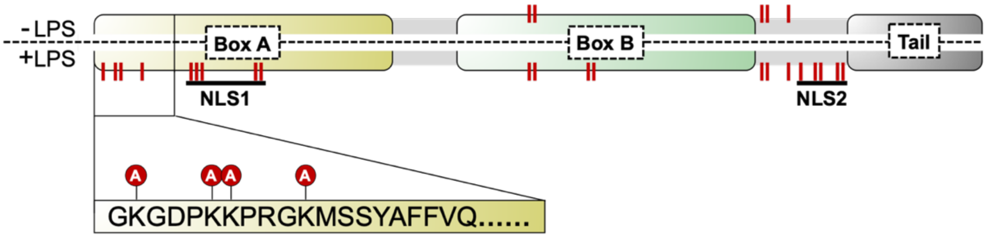


**Figure S1. HMGB1 was hyper-acetylated after LPS stimulation.** The modification sites were highlighted by red lines. Data was adopted from Lu et al.[1]

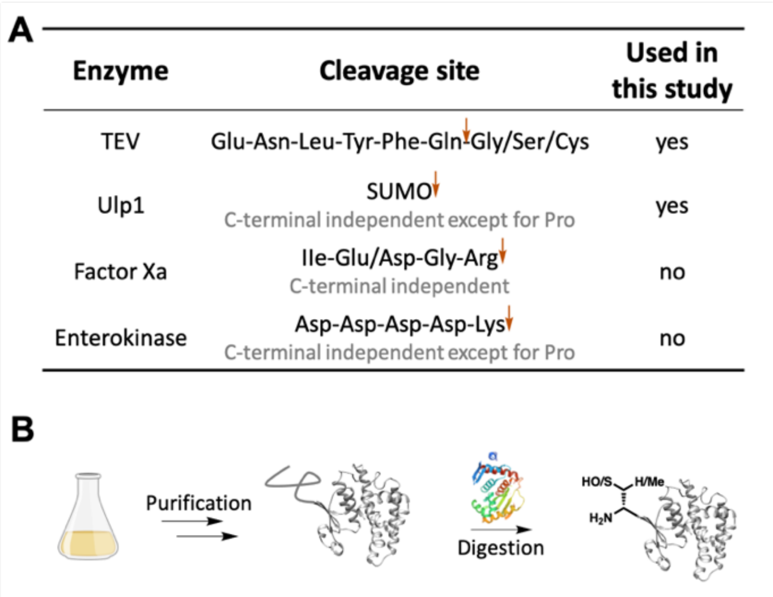

A

| Enzyme | Cleavage site | Used in this study |
|---|---|---|
| TEV | Glu-Asn-Leu-Tyr-Phe-Gln↓Gly/Ser/Cys | yes |
| Ulp1 | SUMO↓ C-terminal independent except for Pro | yes |
| Factor Xa | Ile-Glu/Asp-Gly-Arg↓ C-terminal independent | no |
| Enterokinase | Asp-Asp-Asp-Asp-Lys↓ C-terminal independent except for Pro | no |



**Figure S2. Method to generate recombinant Ser/Thr/Cys N-terminal protein.**

(A). List of enzymes which can produce a Ser/Thr/Cys N-terminal protein.

(B). After Purification, N-terminal Ser/Thr/Cys can be released by indicated enzymes.

**Note**: If the truncated protein is not soluble, we can purify the inclusion body in the denatured buffer (8 M Urea). Then, the protein can be diluted with 2 M Urea. In this concentration of Urea, protein does not precipitate, and enzyme (e.g., TEV, Ulp1) still can cleave the recognition site. After digestion, the protein solution can be further purified by HPLC. Besides, methionine aminopeptidase can remove Met, thus produce a Ser/Thr/Cys N-terminal protein.

| Ser-MBP | 1 | 2 | 3 | 4 | 5 | 6 | 7 | 8 | 9 |
|---|---|---|---|---|---|---|---|---|---|
| Theoretical Mass (Da) | 41066 | 41080 | 42362 | 43173 | 43708 | 43871 | 44456 | 44748 | 45911 |
| Observed Mass (Da) | 41070 (>95%) | 41081 (>95%) | 42363 (75%) | 43178 (50%) | - (<5%) | - (<5%) | 44460 (50%) | - (<5%) | 45914 (75%) |

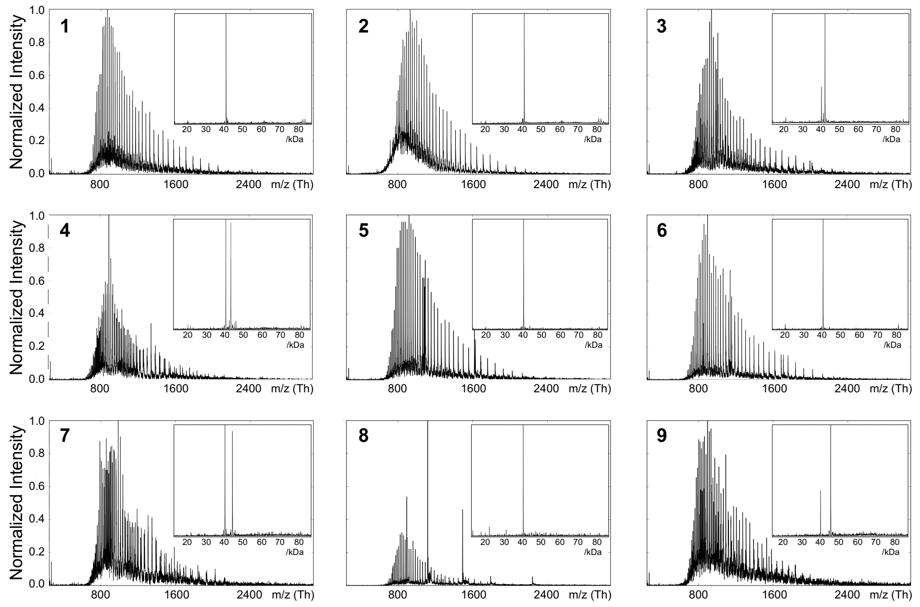


**Figure S3. Mass spectra of ligation products obtained from peptide SAL esters and Ser-MBP.** The conversion rates were calculated according to the deconvolution results.

| Thr-MBP | 1 | 2 | 3 | 4 | 5 | 6 | 7 | 8 | 9 |
|---|---|---|---|---|---|---|---|---|---|
| Theoretical Mass (Da) | 41080 | 41094 | 42376 | 43187 | 43722 | 43885 | 44470 | 44762 | 45925 |
| Observed Mass (Da) | 41088 (>95%) | 41106 (>95%) | 42377 (>95%) | 43191 (85%) | 43719 (<5%) | 43910 (20%) | 44468 (90%) | 44758 (5%) | 45922 (>95%) |

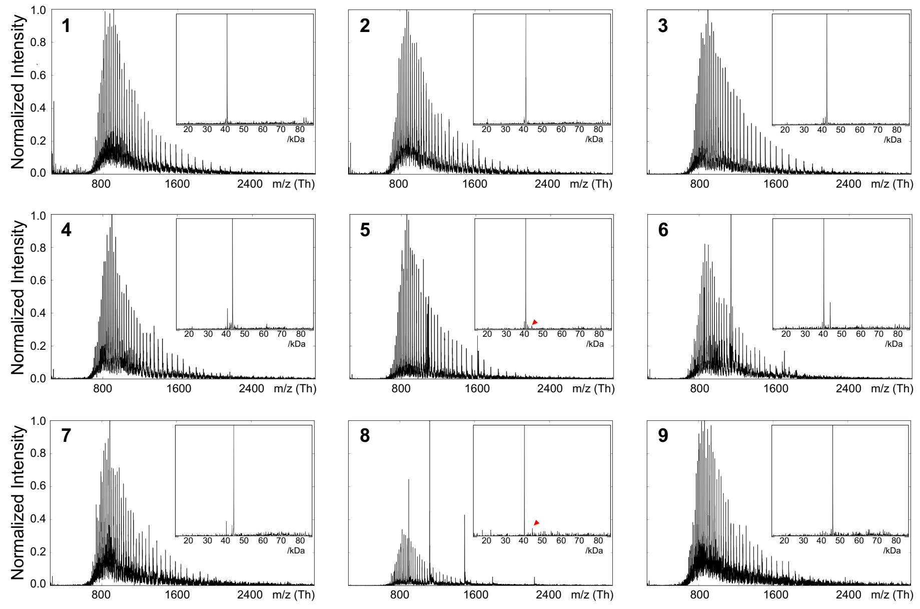


**Figure S4. Mass spectra of ligation products obtained from peptide SAL esters and Thr-MBP.** The conversions were calculated according to the deconvolution results.

| Cys-MBP | 1 | 2 | 3 | 4 | 5 | 6 | 7 | 8 | 9 |
|---|---|---|---|---|---|---|---|---|---|
| Theoretical Mass (Da) | 41082 | 41096 | 42378 | 43189 | 43722 | 43887 | 44472 | 44764 | 45927 |
| Observed Mass (Da) | 41090 (>95%) | 41107 (>95%) | 42381 (75%) | 43194 (70%) | 43723 (>90%) | 43894 (70%) | 44476 (70%) | 44771 (60%) | 45925 (85%) |

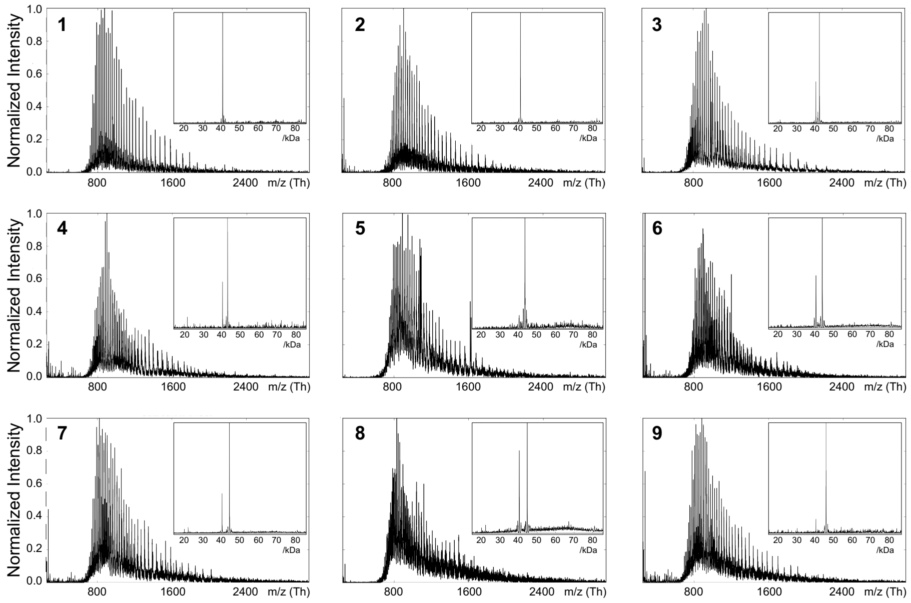


**Figure S5. Mass spectra of ligation products obtained from peptide SAL esters and Cys-MBP.** The conversions were calculated according to the deconvolution results.

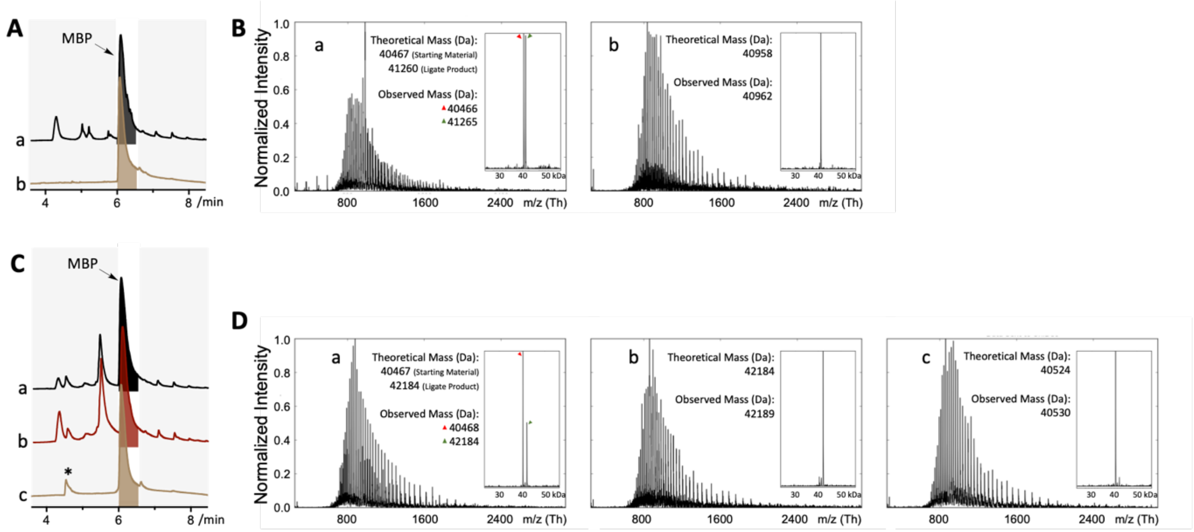


**Figure S6. Purification strategy for STL/CPL mediated protein semi-synthesis.**

(A). HPLC UV traces of reaction mixtures in Disulfide linker mediated purification strategy. a: before purification; b: purified product.

(B). Mass spectra of the products in a and b.

(C). HPLC UV traces of reaction mixtures in His tag and TEV based purification strategy. a: before purification; b: purified product; c: product after TEV digestion.

(D). Mass spectra of products in a, b, and c. *: the remaining peptide SAL ester.

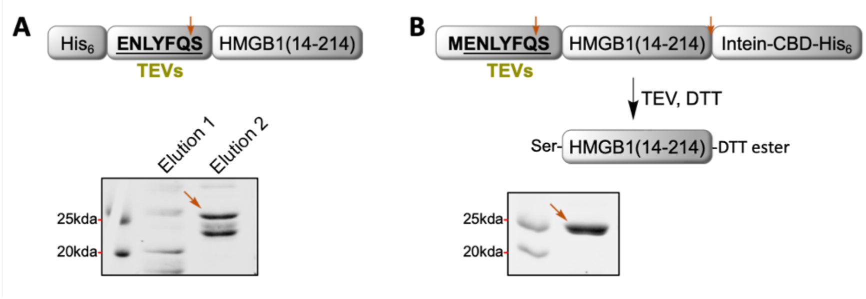


**Figure S7. Purification strategy for HMGB1(13-214).**

(A). The N-terminal His tag purification strategy produced several truncated species of HMGB1.

(B). Intein tag-based purification strategy. Note: DTT ester on C-terminus hydrolyzes slowly.

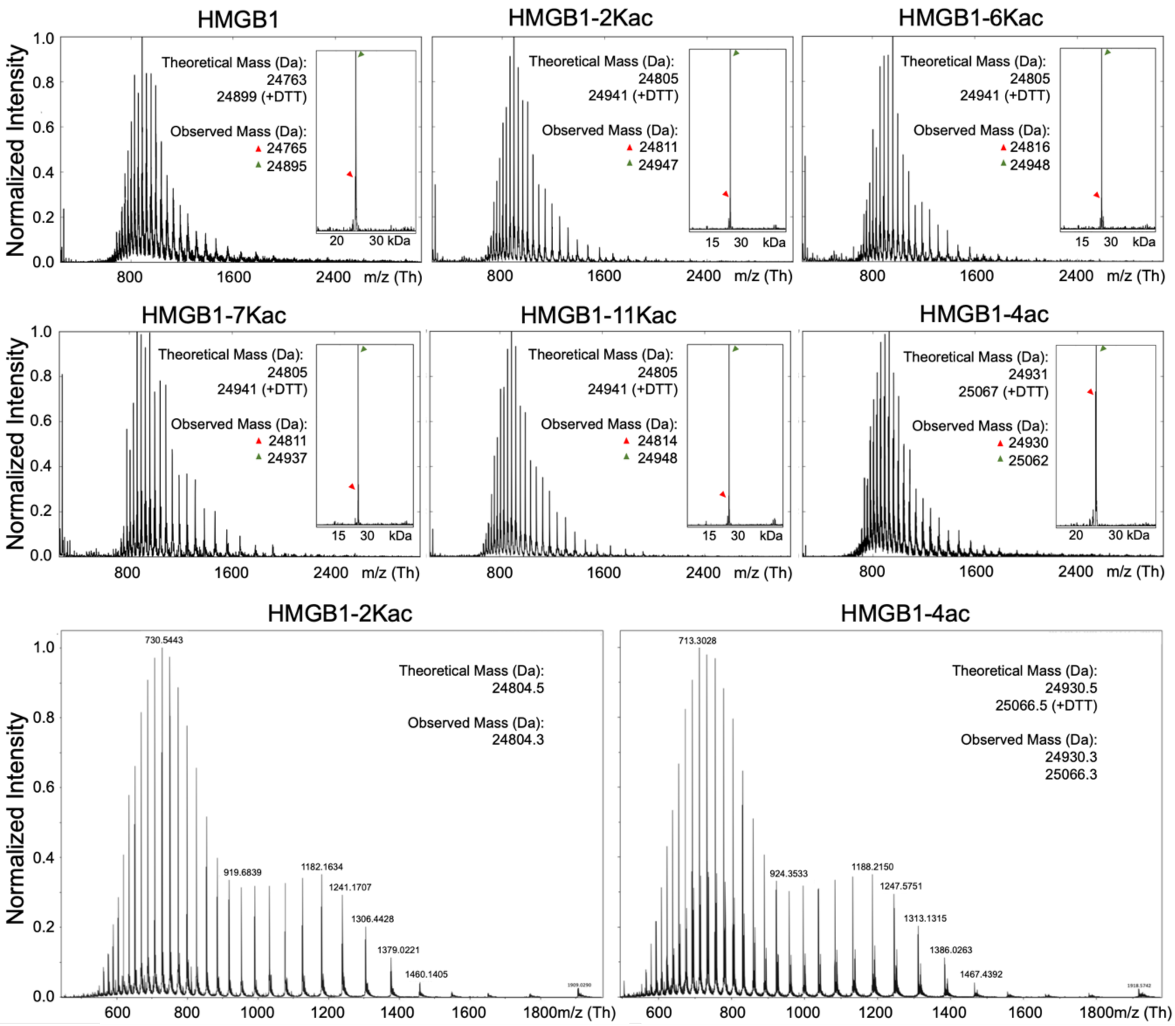


**Figure S8. Mass spectra of synthetic HMGB1 proteins.** The high-resolution mass spectra of HMGB1-2Kac and HMGB1-4ac are also showed. Note: DTT ester on C-terminus hydrolyzed slowly.

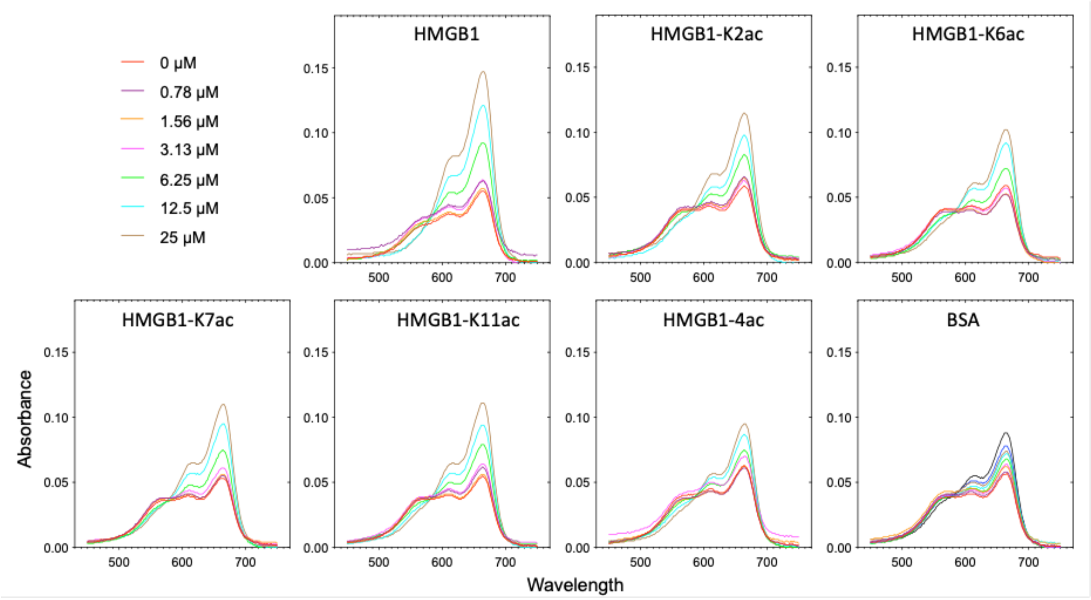


**Figure S9. Absorbance spectra of titration of MB-heparin with different synthetic HMGB1 proteins.**

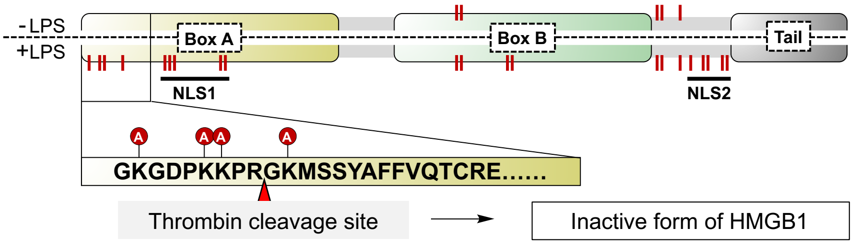


**Figure S10. Thrombin degrades HMGB1 to a less proinflammatory form.**[2]

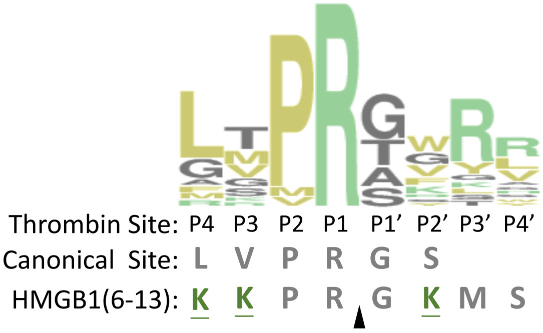


**Figure S11. Position weight matrix for substrate preference of thrombin.**

Raw data was adopted from Gallwitz M et al,[3] in which the recognition preference of thrombin was profiled by phage display technology. They showed the aliphatic residues (e.g. Leu) at P4, Pro at P2, Arg at P1, and small residues at P1' were preferred by thrombin. Positively charged amino acid were coloured as green. Acetylation sites were underlined. The cleavage site is P1^P1'. Canonical site means the thrombin recognition site for recombinant protein production.

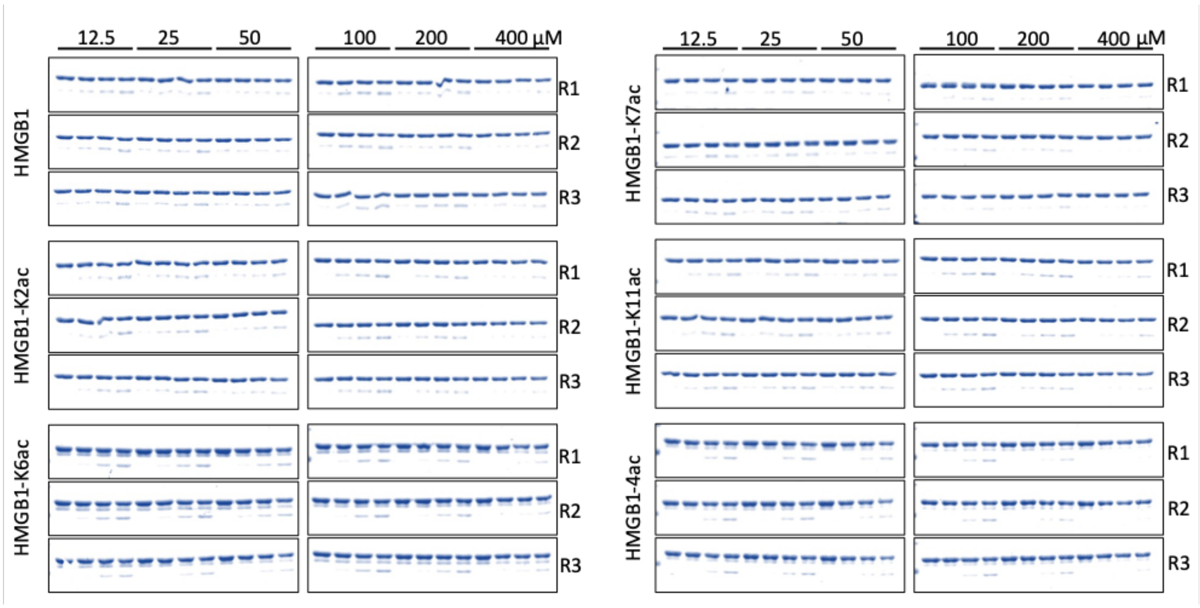


**Figure S12. Original data of SDS-PAGE for Michealis-Menten kinetics analysis.** The bands intensities of full length HMGB1 proteins and cleaved products were quantified by Image lab. Initial reaction velocities were calculated at substrate consumption <25%. Data were fit to a Michaelis–Menten enzyme kinetics model.

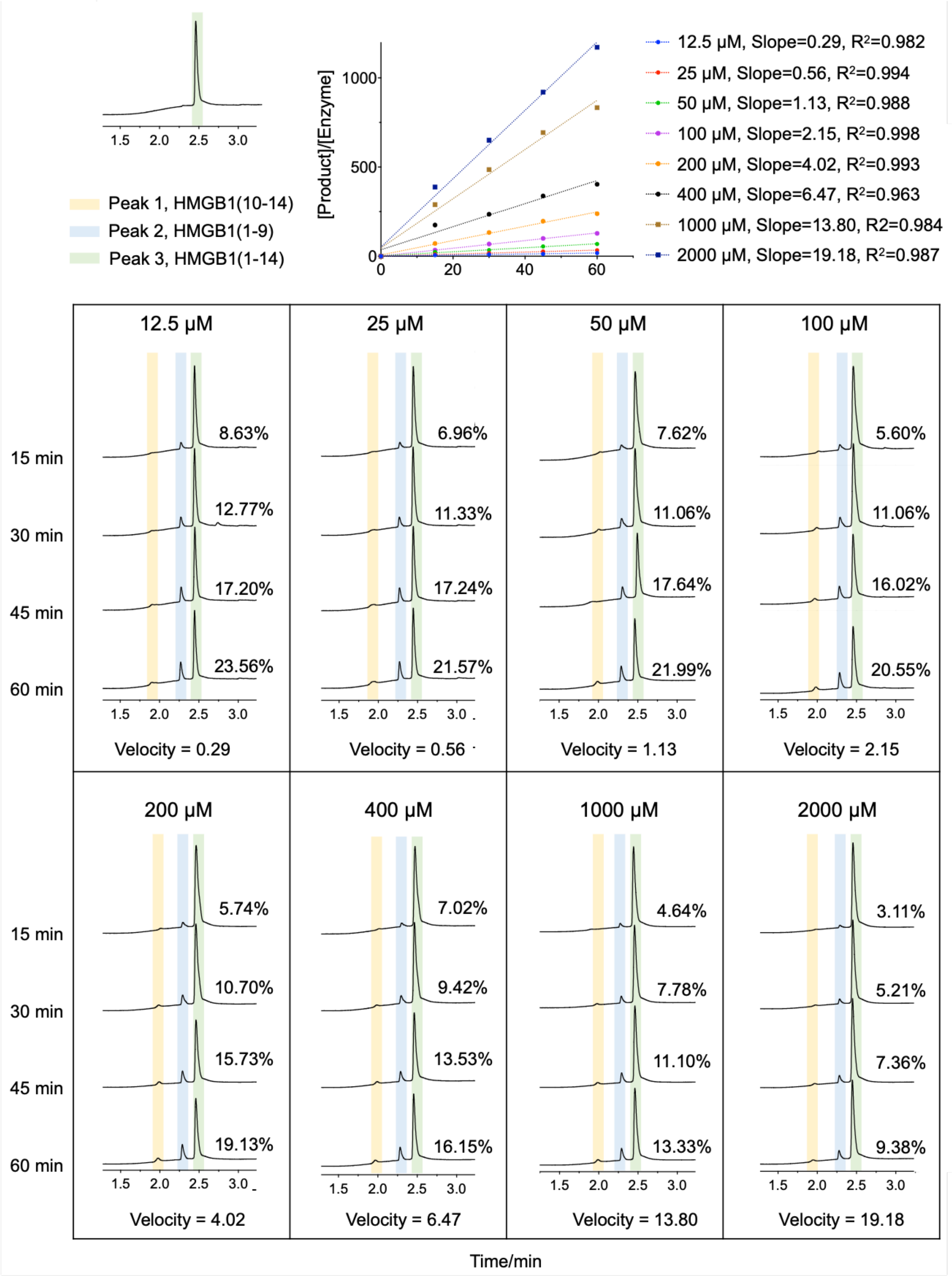


**Figure S13. Representative original data of HPLC for Michealis-Menten kinetics analysis (replicate 1 for HMGB1(1-14)).** The peaks area were integrated Initial reaction velocities were calculated at substrate consumption <25%; consumption = (peak1 + peak2)/(peak1 + peak2 + peak3). Data were fit to a Michaelis–Menten enzyme kinetics model.

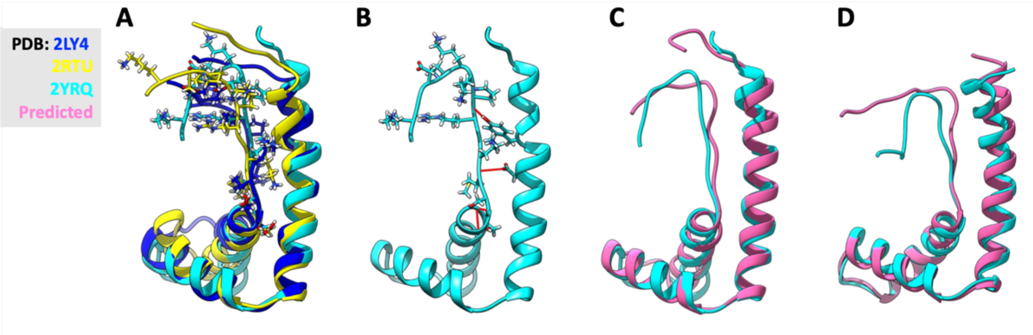


**Figure S14. HMGB1 structures.**

(A). HMGB1 boxA structures which are resolved by independent studies.

(B). Putative hydrogen bonds which could stabilize HMGB1 N-terminal conformation.

(C). HMGB1 boxA structures alignment.

(D). HMGB1 boxB structures alignment. Note: the C-terminal tail and the linker between boxA and boxB are highly flexible.

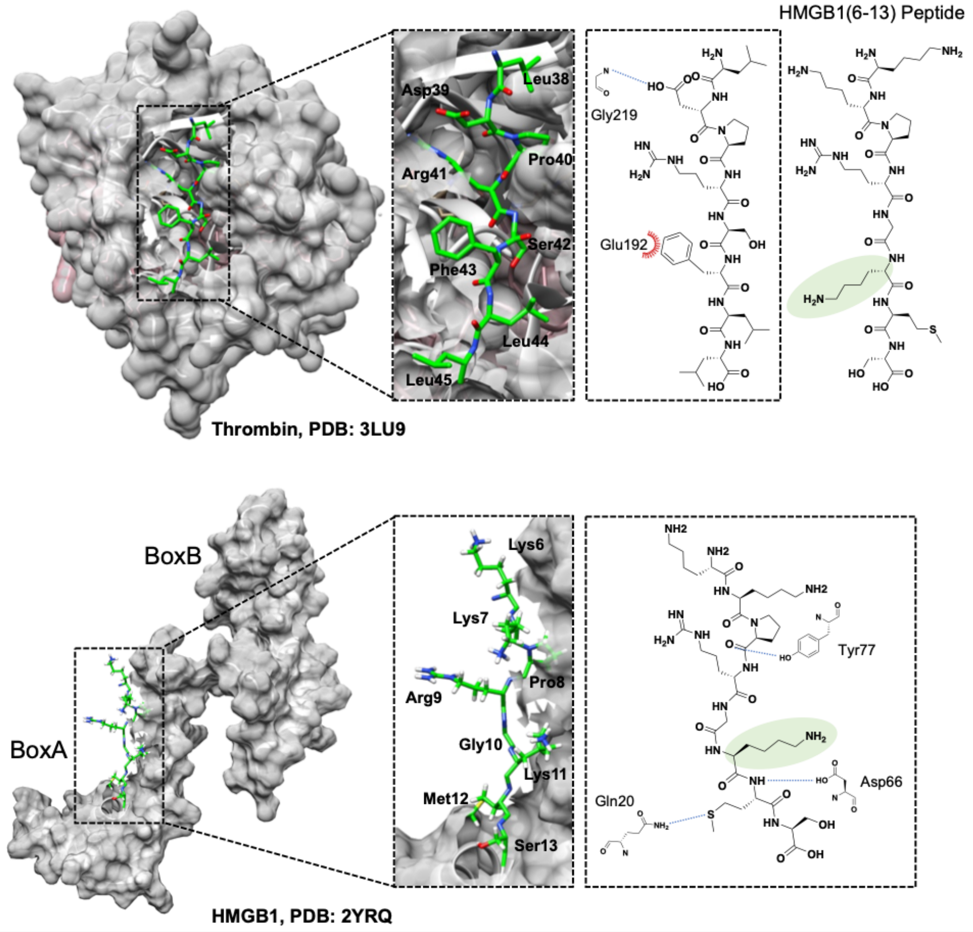


**Figure S15. Structure comparison between thrombin substrate and HMGB1 N-terminus.**

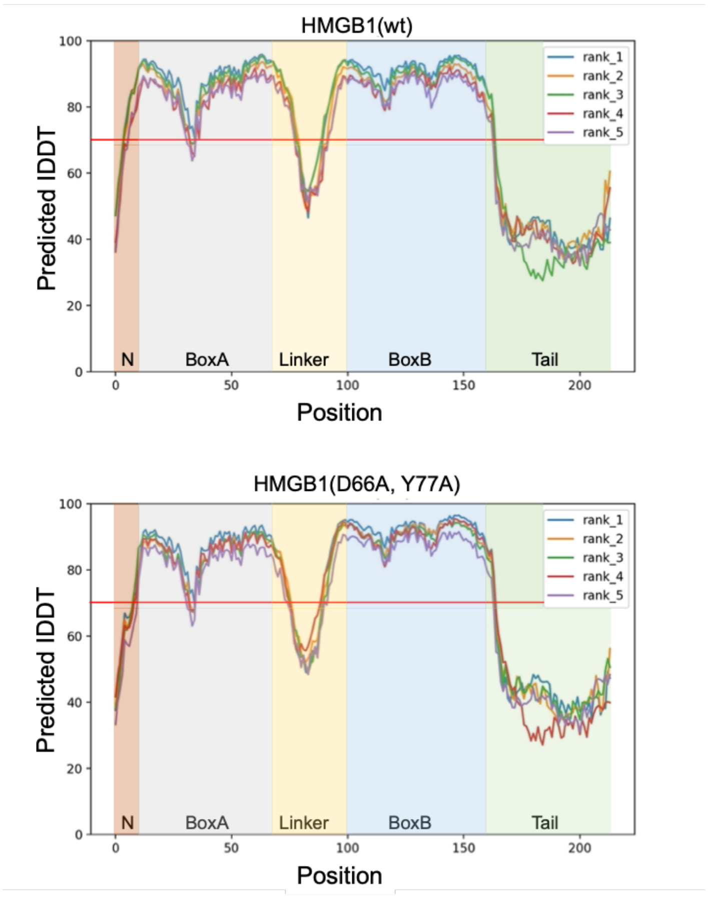


**Figure S16. Predicted local distance difference test (pLDDT) score (per-residue confidence score) for HMGB1(wt) and HMGB1(D66A, Y77A).** Top5 models were plotted. Regions with pLDDT > 90 are expected to be modelled to high accuracy. These should be suitable for any application that benefits from high accuracy; Regions with pLDDT between 70 and 90 are expected to be modelled well (a generally good backbone prediction); Regions with pLDDT between 50 and 70 are low confidence and should be treated with caution.

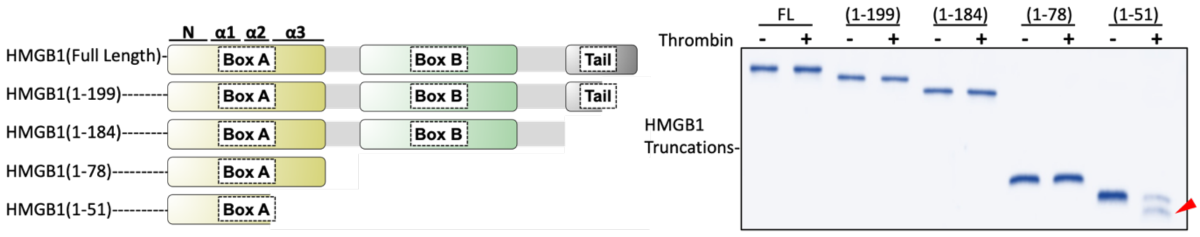


**Figure S17. Thrombin digestion on HMGB1 truncations.**

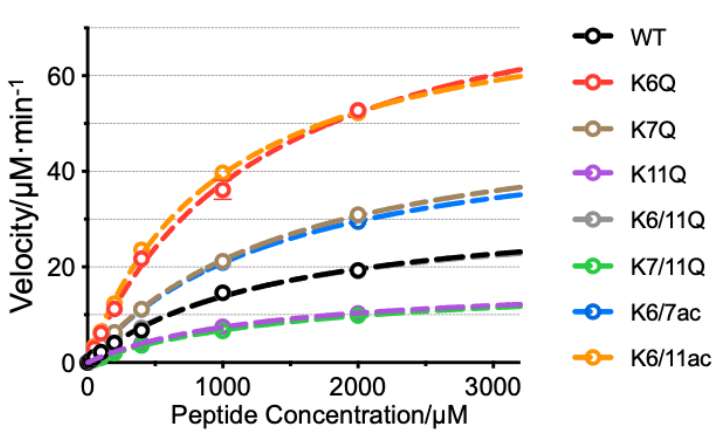


| | | HMGB1 | Vmax (μM·min-1) | Km (mM) | $R^2$ |
|---|---|---|---|---|---|
| Peptide | 1. | HMGB1 | 33.68 ± 2.59 | 1.44 ± 0.21 | 0.9960 |
| | 2. | HMGB1-K6Q | 86.72 ± 6.57 | 1.32 ± 0.19 | 0.9963 |
| | 3. | HMGB1-K7Q | 54.49 ± 4.44 | 1.54 ± 0.21 | 0.9973 |
| | 4. | HMGB1-K11Q | 17.29 ± 0.83 | 1.33 ± 0.12 | 0.9987 |
| | 5. | HMGB1-K6/11Q | 32.32. ± 1.94 | 1.33 ± 0.15 | 0.9976 |
| | 6. | HMGB1-K7/11Q | 17.80 ± 1.67 | 1.64 ± 0.28 | 0.9959 |
| | 7. | HMGB1-K6/7ac | 51.00 ± 1.18 | 1.45 ± 0.06 | 0.9997 |
| | 8. | HMGB1-K6/11ac | 78.39 ± 2.86 | 0.99 ± 0.08 | 0.9987 |

**Figure S18. Michealis-Menten kinetics analysis of thrombin on HMGB1 N-terminal peptides.** Values represent 95% profile likelihood.

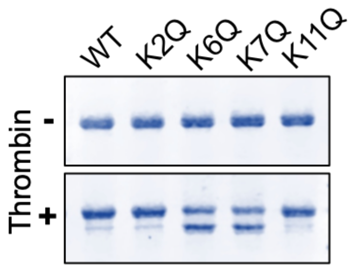


**Figure S19. Thrombin digestion on HMGB1 site mutations.**

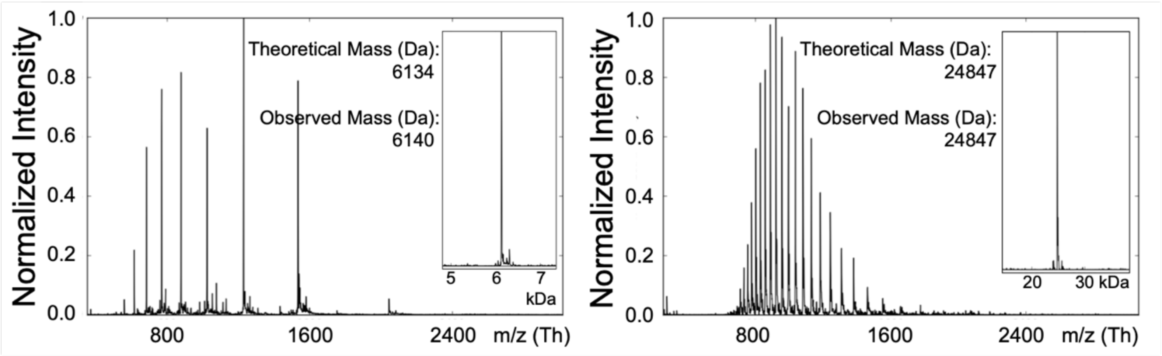


**Figure S20. Mass spectra of the semi-synthetic HMGB1(1-51)-4ac and HMGB1-K6/7ac.**

**A**

| Pyridine/ acetic acid | Remark |
|---|---|
| 1:3 | Can not dissolve proteins at 1 mM. |
| 1:6 | |
| 1:9 | |

**B**

| Pyridine derivatives | Remark |
|---|---|
| Collidine | Can not dissolve proteins at 1 mM. |

**C**

| Co-Solvent | Remark |
|---|---|
| DMF | Can not dissolve proteins at 2 mM. |
| DCM | Can not dissolve proteins at 2 mM. |
| TFE | Can not dissolve proteins at 2 mM. |
| HFIP | Could dissolve all tested proteins. Some salicylaldehyde ester may hydrolyzed quickly. If so, we recommend to use DMSO.<br>Side rection was observed with salicylaldehyde ester( the salicylaldehyde esters were displaced by HFIP). |
| DMSO | Could dissolve all tested proteins. The solution may become viscous during ligation, which may lead the slightly lower ligation conversion than in HFIP.<br>No side rection was observed with salicylaldehyde ester. |

**D**

| Temperature | Remark |
|---|---|
| 25℃ | We always did the the ligation at room temperature. |

**E**

| Equivalent of peptide ester | Remark |
|---|---|
| 2 | All tested peptide ester is soluble. Low conversion (Most reactions are ~20%). |
| 5 | All tested peptide ester is soluble. Medium conversion. |
| 10 | All tested peptide ester is soluble. High conversion. Products were pure. |
| 20 | Some peptide esters is not soluble. High conversion. Some side reactions occurred. |

**Table S1. Condition screening for STL.** To dissolve the protein at 1mM, different ratio of pyridine/acetic acid(A), pyridine derivatives(B) to replace pyridine and co-solvent(C) were screening. We found that protein can be dissolved in HFIP or DMSO at 2 mM firstly then diluted in pyridine/acetic acid for STL. Besides, the equivalent of peptide salicylaldehyde esters was screened(E).

| N-Terminal Amino acid of POI | C- Terminal Amino Acid of Peptide | | |
|---|---|---|---|
| | FAST | SLOW | DECOMPOSED |
| Ser | Ala, Gly, Ser, Gln, Thr, Phe, Cys(SStBu) | Val, Ile, Met, Asn, Tyr, Leu, Trp, Arg, Pro, His | Lys, Asp, Glu |
| Thr | Ala, Gly, Ser, Gln, Thr, Phe, Cys(SStBu) | Val, Ile, Met, Asn, Tyr, Leu, Trp, Arg, Pro, His | |
| Cys | ALL | - | |

**Table S2. Recommendation for disconnected sites for STL/CPL mediated protein semi-synthesis.** Data was original from Liu et al. and Tan et al.[4,5]

#### Synthetic Details

##### 1. General information for reagents and methods

All commercially available amino acids and coupling reagents (purchased from Aldrich and GL Biochem) were used without further purification. All solvents in reagent grade (RCI) or HPLC grade (DUKSAN) were used without purification. Anhydrous dichloromethane (DCM) was freshly distilled from calcium hydride ($CaH_2$) before use. Analytical HPLC was performed on a Waters system equipped with a photodiode array detector (Waters 2996), using a Vydac 218TPTM C18 column (5 μm, 4.6 × 250 mm) at a flow rate of 0.6 mL/min; or on a Waters UPLC H-class system equipped with an ACQUITY UPLC photodiode array detector and a Waters SQ Detector 2 mass spectrometer using a Waters ACQUITY BEH C18 column (1.7 μm, 130 Å, 2.1 × 50 mm) at a flow rate of 0.4 mL/min. Preparative HPLC was performed on a Waters system, using a Vydac 218TPTM C18 column (10 μm, 22 × 250 mm) at a flow rate of 10 mL/min or a Vydac 218TPTM C18 column (10 μm, 30 × 250 mm) at a flow rate of 20 mL/min. Mobile phases of HPLC used are as followed: Solvent A: 0.1% TFA (v/v) in acetonitrile ($CH_3CN$, ACN); Solvent B: 0.1% TFA (v/v) in water. Mass analysis were performed with a Waters 3100 mass spectrometer.

##### 2. General experimental procedures

##### 2.1 Solid-phase peptide synthesis (SPPS)

The solid phase peptide synthesis was carried out manually using 2-chloro-trityl resin (GL Biochem, loading capacity: 0.5 mmol/g). 2-Chloro-trityl chloride resin was swollen in anhydrous $CH_2Cl_2$ for 30 min and then it washed with $CH_2Cl_2$ (5 mL × 3). After that, a solution of Fmoc-Xaa-OH (4.0 equiv. relative to resin loading capacity) and DIEA (8.0 equiv. relative to resin capacity) in $CH_2Cl_2$ was added and the resin was shaken at room temperature for 2 h to load the first amino acid. Then the resin was washed with DMF (5 mL × 3) and $CH_2Cl_2$ (5 mL × 3), and subsequently treated with a solution of $CH_2Cl_2$/$CH_3OH$/DIEA (17:2:1, v/v/v, 5 mL) for 1 h for capping. The resin was washed with DMF (5 mL × 3), $CH_2Cl_2$ (5 mL × 3), and DMF (5 mL × 3). Finally, it was subjected to iterative peptide assembly (Fmoc-SPPS). The deFmoc solution was the mixture of piperidine/DMF 20/80 (v/v). For the deFmoc step, the resin was treated with deFmoc solution at R.T. for 20 min. The deFmoc solution was removed, then the resin was washed with DMF (5 mL × 3), $CH_2Cl_2$ (5 mL × 3), and DMF (5 mL × 3).

For the coupling step, a solution of Fmoc protected amino acid or Boc protected amino acid (4.0 equiv. according to the resin capacity), HATU (4.0 equiv.) and DIEA (10 equiv.) in DMF was gently agitated with the resin at room temperature for 1h. Double coupling was employed for coupling Histidine. The resin was washed with DMF (5 mL × 3), $CH_2Cl_2$ (5 mL × 3), and DMF (5 mL × 3). The following Fmoc amino acids and Boc amino acids from GL Biochem were employed: Fmoc-Ala-OH, Fmoc-Cys(Acm)-OH, Fmoc-Cys(Trt)-OH Fmoc-Asp(OtBu)-OH, Fmoc-Glu(OtBu)-OH, Fmoc-Phe-OH, Fmoc-Gly-OH, Fmoc-His(Trt)-OH, Fmoc-His(Boc)-OH, Fmoc-Ile-OH, Fmoc-Lys(Boc)-OH, Fmoc-Leu-OH, Fmoc-Met-OH, Fmoc-Asn(Trt)-OH, Fmoc-Pro-OH, Fmoc-Gln(Trt)-OH, Fmoc-Arg(Pbf)-OH, Fmoc-Ser(tBu)-COOH, Fmoc-Thr(tBu)-COOH, Fmoc-Val-OH, Fmoc-Trp(Boc)-OH, Fmoc-Tyr(tBu)-OH, Boc-Ala-OH, Boc-Met-OH and Boc-Cys(StBu)-OH.

#### 2.2 Cleavage fully protected peptide from 2-chloro-trityl chloride resin

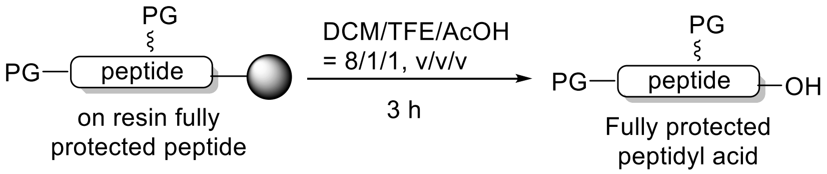


The on-resin fully protected peptide, obtained as described in the **General Experimental Procedures 2.1**, was subjected to the mild acidic cleavage cocktail (5-10 mL) of $CH_2Cl_2$/AcOH/trifluoroethanol (8/1/1, v/v/v), 3 times for 60 min each. Following filtration, the resulting cleavage solutions were combined and concentrated to afford the crude protected peptide with the free carboxylic acid at the *C*-terminus.

#### 2.3 Synthesis of model C-terminal peptide SAL esters

##### 2.3.1 Direct coupling for preparation of C-terminal Gly and Pro peptide SAL esters:

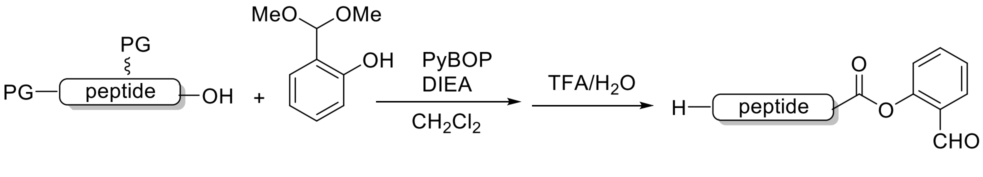


Fully protected crude peptide (1.0 equiv.) obtained from **General Experimental Procedures 2.2** was dissolved in dry DCM at a concentration of 10 mM. DIEA (6.0 equiv.) and benzotriazol-1-yl-oxytripyrrolidinophosphonium hexafluorophosphate (PyBOP) (3.0 equiv.) were added, followed by salicylaldehyde dimethyl acetal (30.0 equiv.). The reaction mixture was stirred at room temperature for overnight. After that, the solvent was removed under reduced pressure and the resulting residue was treated with TFA/$H_2O$ (95:5, v/v). After global deprotection for 2 h, TFA was blown off and the oily residue was triturated with diethyl ether and centrifuged. The precipitate was pelleted and the ether was subsequently decanted. The resulting solid was purified by HPLC and lyophilized to give the peptide SAL esters as a white solid.

##### 2.3.2 "N+1" strategy for the preparation of C-terminal Ser, Met, Ala, Phe, Val, Leu, Ile, and Thr peptide SAL esters:

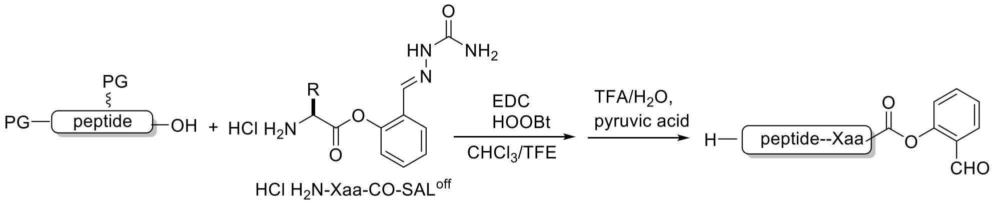


The fully protected peptidyl acid (1.0 equiv.) obtained from **General Experimental Procedures 2.2** was dissolved in $CHCl_3$/trifluoroethanol (10 mM, 3/1, v/v), then the corresponding L-Amino acid derived salicylaldehyde semicarbazone ester hydrochloride HCl·$H_2N$-Xaa-CO-$SAL^{off}$ (6.0 equiv.), synthesized according to the procedure[6] and hydroxy-3,4-dihydro-4-oxo-1,2,3-benzotriazine (HOOBt) (3.0 equiv.) were added. Finally, N-(3-dimethylaminopropyl)-N'-ethylcarbodiimide (EDC) (3.0 equiv.) was added. The reaction mixture was stirred for 3 h to form the crude protected C-terminal peptide $SAL^{off}$ ester. After that, the solvent was removed under reduced pressure and the resulting residue was treated with TFA/$H_2O$ (95:5, v/v) containing pyruvic acid (100 equiv.) for 3 h. After that, TFA was blown off and the oily residue was triturated with diethyl ether and centrifuged. The precipitate was pelleted and the ether was subsequently decanted. The resulting solid was purified by HPLC and lyophilized to give the peptide SAL esters as a white solid.

#### 3. Synthesis of LSQRGG-CO-SAL ester

The H-LSQRGG-CO-SAL was synthesized according to the **General Experimental Procedures 2.3.1**.

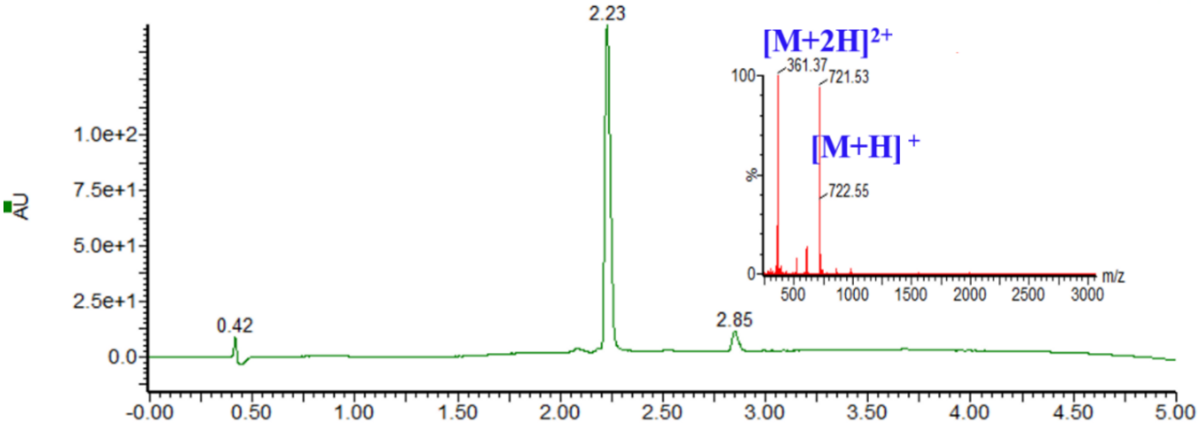


UV trace and corresponding MS from LC-MS analysis of purified H-LSQRGG-CO-SAL. ESI-MS calcd. molecular weight: 720.79. $[M+H]^{+}$ m/z = 721.79, $[M+2H]^{2+}$ m/z = 361.39, found 721.53, 361.37.

**4. Synthesis of H-LSQRGA-CO-SAL ester**

The H-LSQRGA-CO-SAL was synthesized according to the **General Experimental Procedures 2.3.2**.

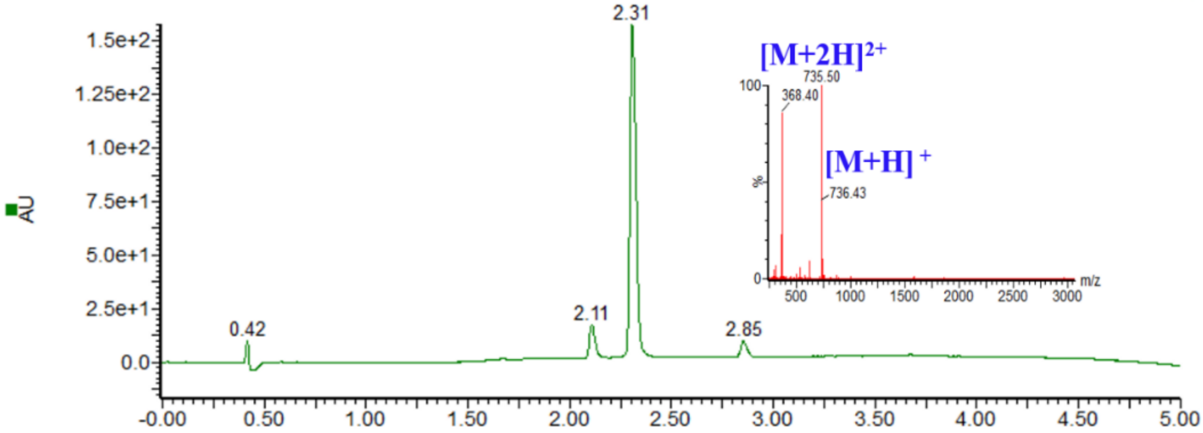


UV trace and corresponding MS from LC-MS analysis of purified H-LSQRGA-CO-SAL. ESI-MS calcd. molecular weight: 734.81. $[M+H]^{+}$ m/z = 735.81, $[M+2H]^{2+}$ m/z = 368.40, found 735.50, 368.40.

**5. Synthesis of Ac-ETTTQGPGVLLPLPKGAC(StBu)-CO-SAL ester**

The Ac-ET TTQGPGVLLP LPKGAC(StBu)-CO-SAL was synthesized according to the **General Experimental Procedures 2.3.2**.

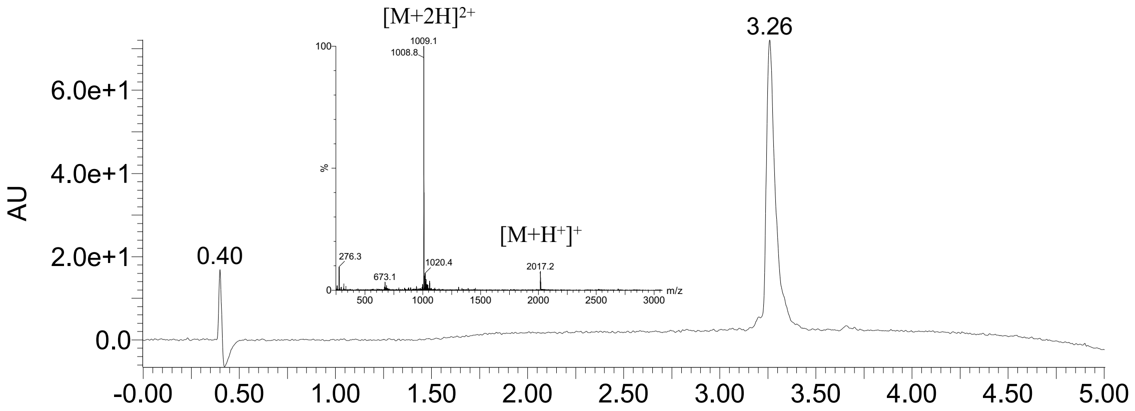


UV trace and corresponding MS from LC-MS analysis of purified product. ESI-MS calcd. molecular weight: 2016.40. $[M+H]^+$ m/z = 2017.40, $[M+2H]^{2+}$ m/z = 1009.20, found 2017.20, 1009.10.

#### 6. Synthesis of Biotin-C(Acm)SRAARGTIGARRTGQPLKEDPS-CO-SAL ester

The Biotin-C(Acm)SRAARGTIGARRTGQPLKEDPS-CO-SAL was synthesized according to the **General Experimental Procedures 2.3.2.**

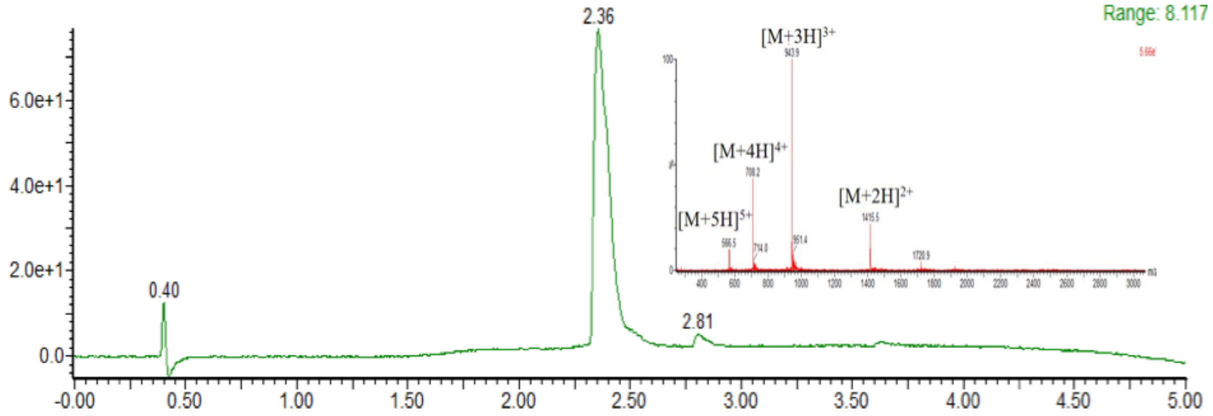


UV trace and corresponding MS from LC-MS analysis of purified product. ESI-MS calcd. molecular weight: 2828.21. $[M+2H]^{2+}$ m/z = 1415.6, $[M+3H]^{3+}$ m/z = 944.0, $[M+4H]^{4+}$ m/z = 708.3, $[M+5H]^{5+}$ m/z = 566.8, found 1415.5, 943.9, 708.2, 566.5.

#### 7. Synthesis of Fmoc-HN-TLAEAQTETC(Acm)TVAPRERQNC(StBu)GFPGVTP-CO-SAL ester

The Fmoc-HN-TLAEAQTETC(Acm)TVAPRERQNC(StBu)GFPGVTP-CO-SAL was synthesized according to the **General Experimental Procedures 2.3.1.**

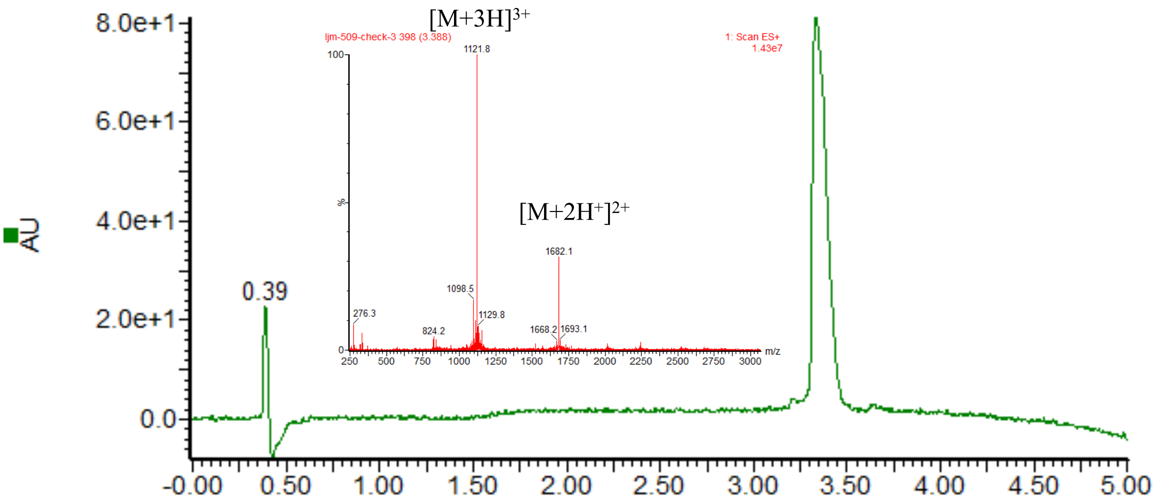


UV trace and corresponding MS from LC-MS analysis of purified product. ESI-MS calcd. molecular weight: 3362.80. $[M+2H]^{2+}$ m/z = 1682.4, $[M+3H]^{3+}$ m/z = 1121.93, found 1682.1, 1121.8.

**8. Synthesis of Fmoc-SEAVLRGQALLVKSSQPWEPLQLHVDKAV-CO-SAL ester**

The Fmoc-SEAVLRGQALLVKSSQPWEPLQLHVDKAV-CO-SAL was synthesized according to the **General Experimental Procedures 2.3.2.**

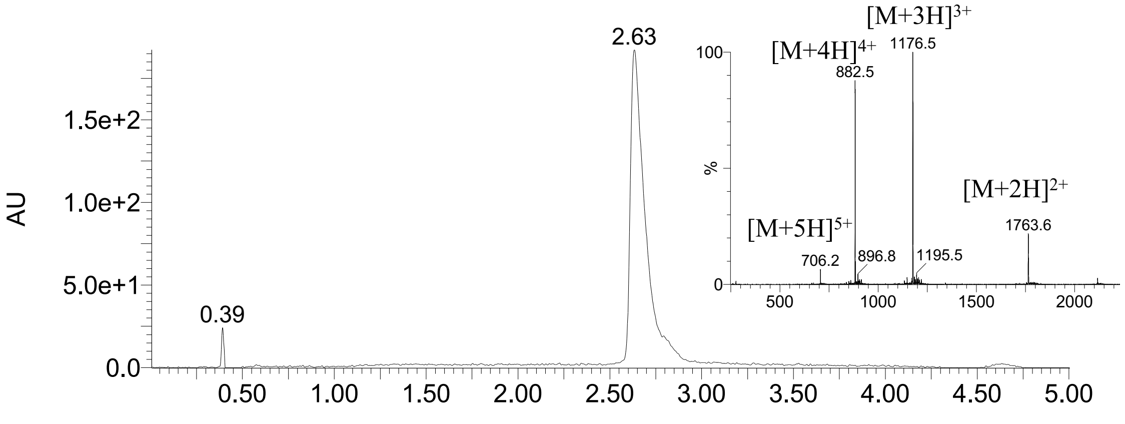


UV trace and corresponding MS from LC-MS analysis of purified product. ESI-MS calcd. molecular weight: 3526.06. $[M+2H]^{2+}$ m/z = 1764.03, $[M+3H]^{3+}$ m/z = 1176.35, $[M+4H]^{4+}$ m/z = 882.51, $[M+5H]^{5+}$ m/z = 706.21, found 1763.6, 1176.5, 882.5, 706.2.

**9. Synthesis of Ac-SESSSKSSQPLASKQEKDGTEKRGRGRPRKQPPVSPG-CO-SAL ester**

The Ac-SESSSKSSQPLASKQEKDGTEKRGRGRPRKQPPVSPG-CO-SAL was synthesized according to the **General Experimental Procedures 2.3.1.**

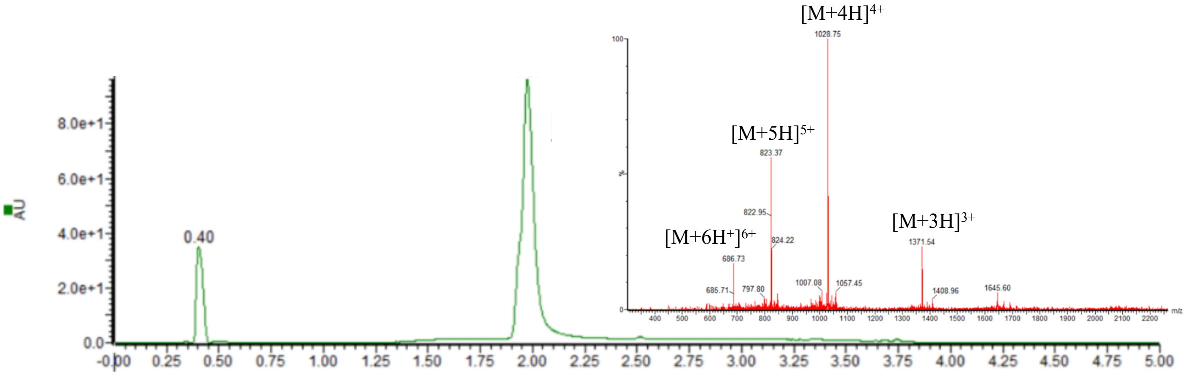


UV trace and corresponding MS from LC-MS analysis of purified product. ESI-MS calcd. molecular weight: 4111.51. $[M+3H]^{3+}$ m/z = 1371.50, $[M+4H]^{4+}$ m/z = 1028.89, $[M+5H]^{5+}$ m/z = 823.30, $[M+6H]^{6+}$ m/z = 686.25, found: 1371.54, 1028.75, 823.37, 686.73.

#### 10. Synthesis of Fmoc-HN-TGANRDLELPWLEQQGPASHHRRQLGPQGPPHLVADP-CO-SAL ester

The Fmoc-HN-TGANRDLELPWLEQQGPASHHRRQLGPQGPPHLVADP-CO-SAL was synthesized according to the **General Experimental Procedures 2.3.1.**

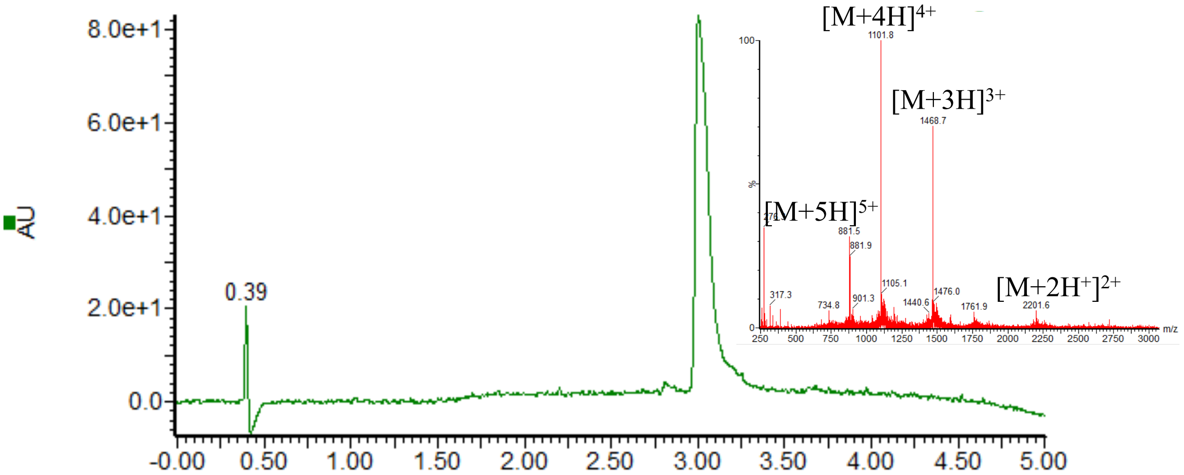


UV trace and corresponding MS from LC-MS analysis of purified product. ESI-MS calcd. molecular weight: 4402.87. $[M+2H]^{2+}$ m/z = 2202.44, $[M+3H]^{3+}$ m/z = 1468.62, $[M+4H]^{4+}$ m/z = 1101.72, $[M+5H]^{5+}$ m/z = 881.57, found 2201.6, 1468.7, 1101.8, 881.5.

#### 11. Synthesis of Biotin-WRRKRKEKQSETSPKEFLTIYEDVKDLKTRRNHEQEQTF PGGG-CO-SAL ester

The Biotin-WRRKRKEKQSETSPKEFLTIYEDVKDLKTRRNHEQEQTFPGGG-OSAL was synthesized according to the **General Experimental Procedures 2.3.1.**

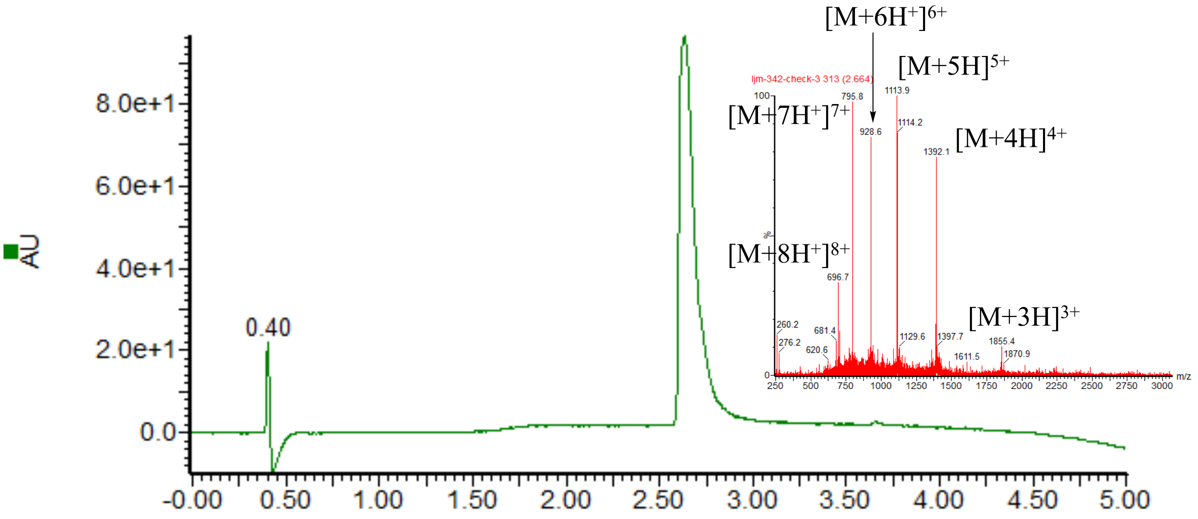


UV trace and corresponding MS from LC-MS analysis of purified product. ESI-MS calcd. molecular weight: 5565.68. $[M+3H]^{3+}$ m/z = 1856.23, $[M+4H]^{4+}$ m/z = 1392.42, $[M+5H]^{5+}$ m/z = 1114.14, $[M+6H]^{6+}$ m/z = 928.61, $[M+7H]^{7+}$ m/z = 796.10, $[M+8H]^{8+}$ m/z = 696.71, found 1855.4, 1392.1, 1113.9, 928.6, 795.8, 696.7.

#### 12. Synthesis of disulfide-containing peptide SAL ester

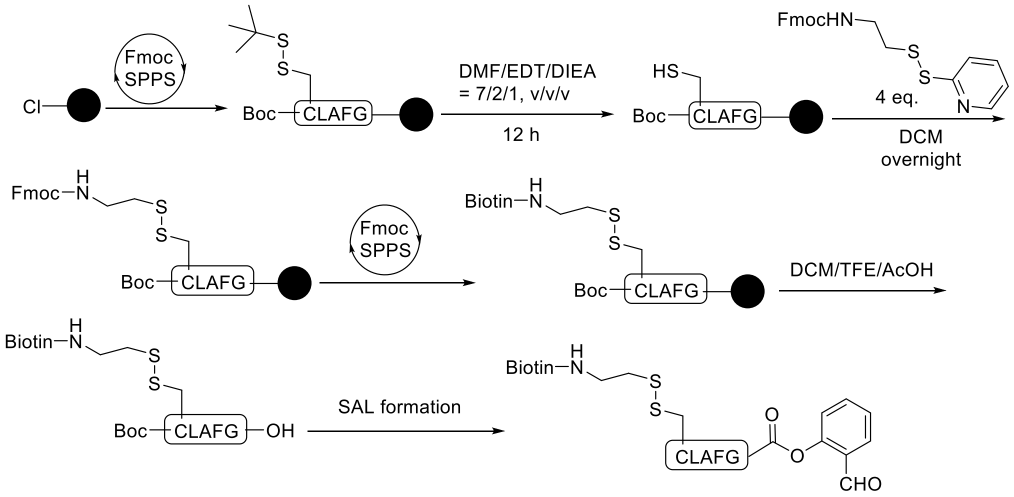


The on resin fully protected peptide obtained using **General Experimental Procedures 2.1** was treated with 5 mL EDT/DIEA/DMF (2/1/7, v/v/v) and shaken for 12 h at room temperature to selectively remove the *tert*-butylthio (StBu) protecting group. Then, the resin was washed with DMF (5 mL × 3) and $CH_2Cl_2$ (5 mL × 3), and subsequently treated with a solution of (9H-fluoren-9-yl)methyl 2-(pyridin-2-yldisulfanyl)ethylcarbamate (synthesized according to reported procedures)[7] (4.0 equiv. relative to resin loading) in DCM for 12 h. After that, the resin was washed with DMF (5 mL × 3), $CH_2Cl_2$ (5 mL × 3), and DMF (5 mL × 3). Subsequently, the Fmoc group on the peptide was removed by treating the resin with a mixture of piperidine/DMF 20/80 (v/v) (4 mL) for 20 min and washed with DMF (5 mL × 3), $CH_2Cl_2$ (5 mL × 3), and DMF (5 mL × 3). Biotin was attached by coupling under standard condition in the next step. Finally, the SAL ester product was prepared according to the **General Experimental Procedures 2.3.1.** The crude peptide SAL ester was purified by preparative reverse-phase HPLC (20-60% $CH_3CN/H_2O$ over 30 min) and lyophilized to afford the desired SAL ester (6 mg, 3.4% yield).

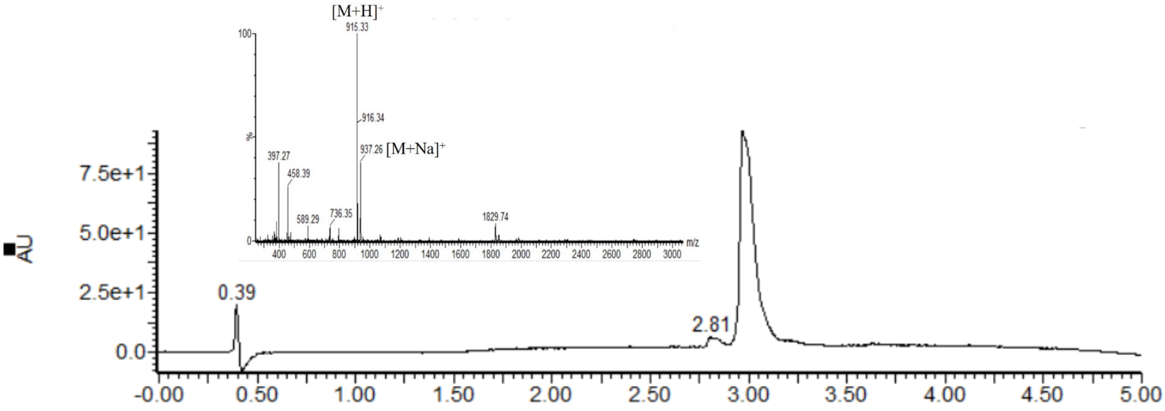


UV trace and corresponding MS from LC-MS analysis of purified product. ESI-MS calcd. molecular weight: 915.15. $[M+H]^+$ m/z = 915.34, $[M+Na]^+$ m/z = 937.34, found 915.33, 937.26.

#### 13. Synthesis of Ac-HHHHHHENLYFQG-CO-SAL ester

The Ac-HHHHHHENLYFQG-CO-SAL was synthesized according to the **General Experimental Procedures 2.3.1.**

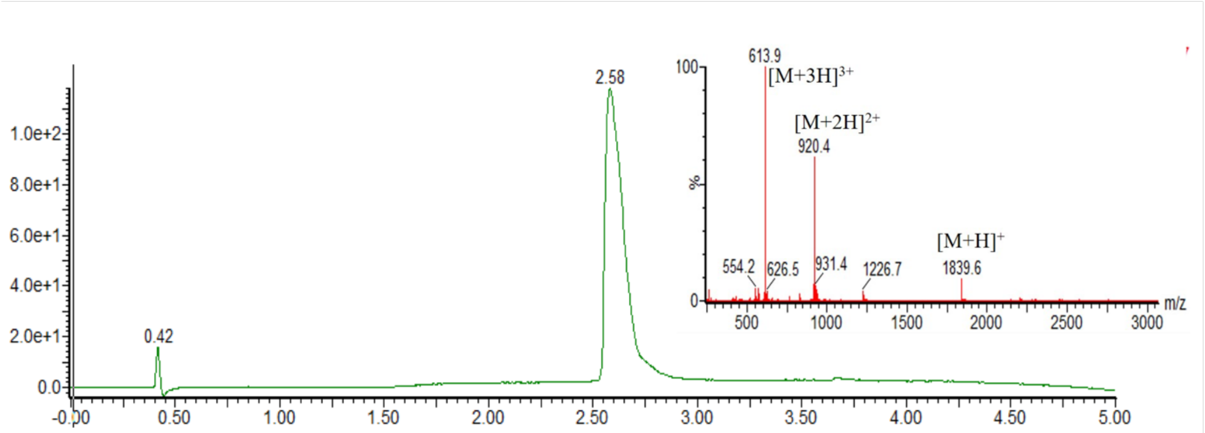


UV trace and corresponding MS from LC-MS analysis of purified product. ESI-MS calcd. molecular weight: 1838.93. $[M+H]^+$ m/z = 1839.93 $[M+2H]^{2+}$ m/z = 920.47, $[M+3H]^{3+}$ m/z = 613.98, found 1839.6, 920.4, 613.9.

#### 14. Synthesis of Ac-HHHHHHENLYFQGKGDPKKPRGKM-CO-SAL ester

The Ac-HHHHHHENLYFQGKGDPKKPRGKM-CO-SAL was synthesized according to the **General Experimental Procedures 2.3.2.**

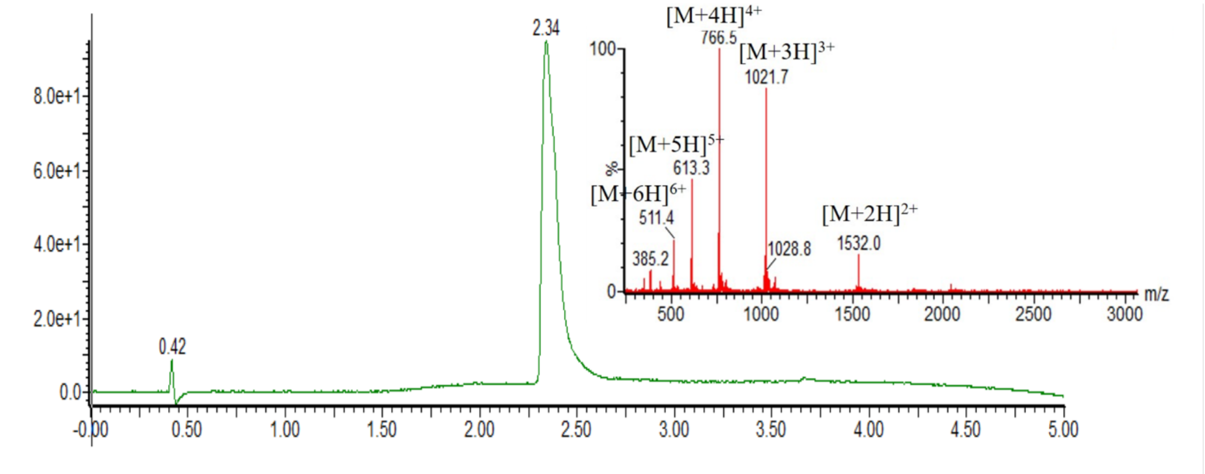


UV trace and corresponding MS from LC-MS analysis of purified product. ESI-MS calcd. molecular weight: 3062.43. $[M+2H]^{2+}$ m/z = 1532.2, $[M+3H]^{3+}$ m/z = 1021.8, $[M+4H]^{4+}$ m/z = 766.6, $[M+5H]^{5+}$ m/z = 613.4, $[M+6H]^{6+}$ m/z = 511.4, found 1532.0, 1021.7, 766.5, 613.3, 511.4.

#### 15. Synthesis of Ac-HHHHHHENLYFQGK(ac)GDPKKPRGKM-CO-SAL ester

The Ac-HHHHHHENLYFQGK(ac)GDPKKPRGKM-CO-SAL was synthesized according to the **General Experimental Procedures 2.3.2.**

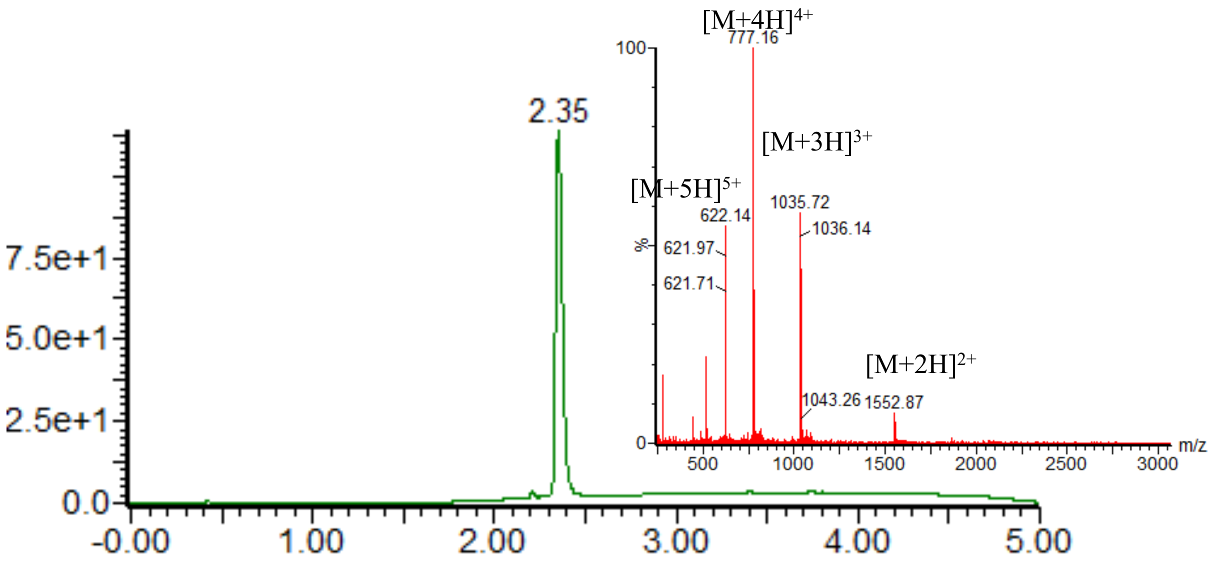


UV trace and corresponding MS from LC-MS analysis of purified product. ESI-MS calcd. molecular weight: 3104.47. $[M+2H]^{2+}$ m/z = 1553.2, $[M+3H]^{3+}$ m/z = 1035.8, $[M+4H]^{4+}$ m/z = 777.1, $[M+5H]^{5+}$ m/z = 621.9, found 1552.9, 1035.7, 777.2, 622.1.

**16. Synthesis of Ac-HHHHHHENLYFQGKGDPK(ac)KPRGKM-CO-SAL ester**

The Fmoc-Lys(Ac)-OH was synthesized according to the reported procedure.[8] The Ac-HHHHHHENLYFQGKGDPK(ac)KPRGKM-CO-SAL was synthesized according to the **General Experimental Procedures 2.3.2.**

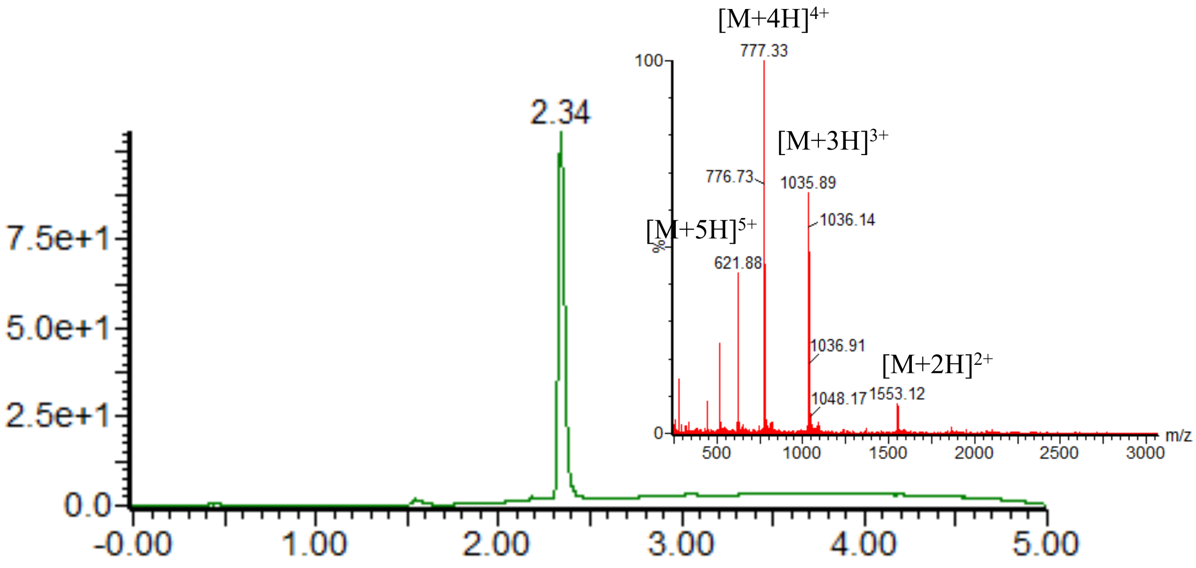


UV trace and corresponding MS from LC-MS analysis of purified product. ESI-MS calcd. molecular weight: 3104.47. $[M+2H]^{2+}$ m/z = 1553.2, $[M+3H]^{3+}$ m/z = 1035.8, $[M+4H]^{4+}$ m/z = 777.1, $[M+5H]^{5+}$ m/z = 621.9, found 1553.1, 1035.9, 777.3, 621.9.

**17. Synthesis of Ac-HHHHHHENLYFQGKGDPKK(ac)PRGKM-CO-SAL ester**

The Ac-HHHHHHENLYFQGKGDPKK(ac)PRGKM-CO-SAL was synthesized according to the **General Experimental Procedures 2.3.2.**

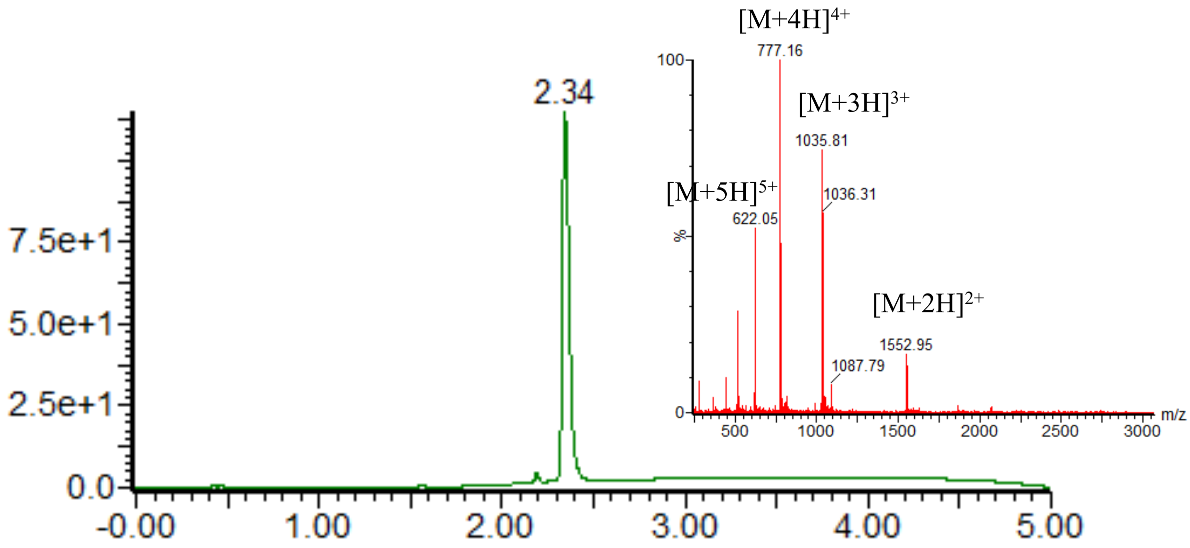


UV trace and corresponding MS from LC-MS analysis of purified product. ESI-MS calcd. molecular weight: 3104.47. $[M+2H]^{2+}$ m/z = 1553.2, $[M+3H]^{3+}$ m/z = 1035.8, $[M+4H]^{4+}$ m/z = 777.1, $[M+5H]^{5+}$ m/z = 621.9, found 1553.0, 1035.8, 777.2, 622.0.

**18. Synthesis of Ac-HHHHHHENLYFQGKGDPKKPRGK(ac)M-CO-SAL ester**

The Ac-HHHHHHENLYFQGKGDPKKPRGK(ac)M-CO-SAL was synthesized according to the **General Experimental Procedures 2.3.2.**

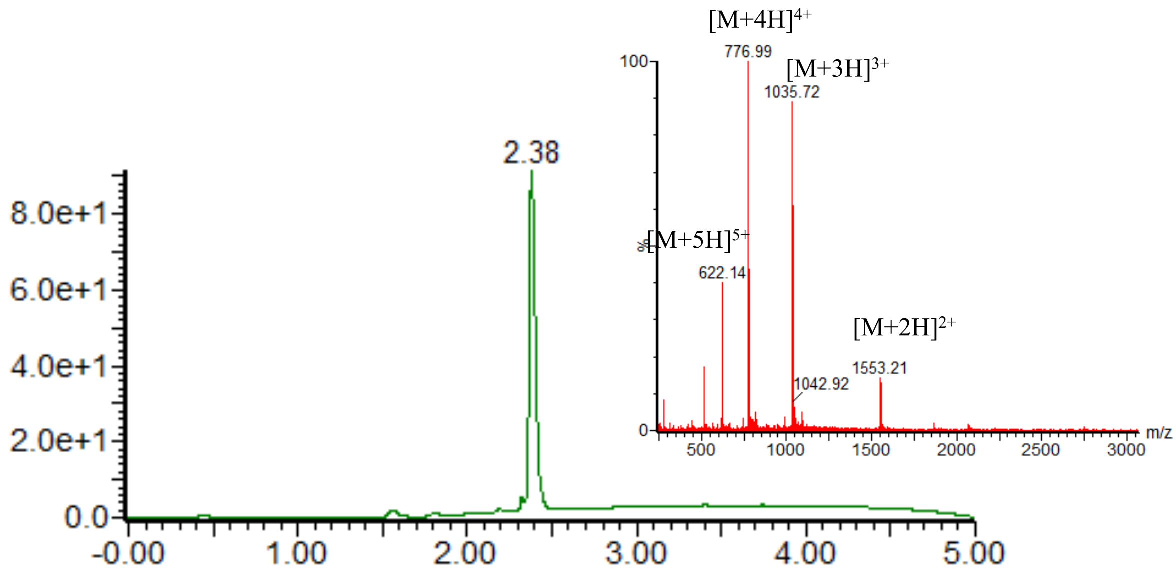


UV trace and corresponding MS from LC-MS analysis of purified product. ESI-MS calcd. molecular weight: 3104.47. $[M+2H]^{2+}$ m/z = 1553.2, $[M+3H]^{3+}$ m/z = 1035.8, $[M+4H]^{4+}$ m/z

= 777.1, $[M+5H]^{5+}$ m/z = 621.9, found 1553.2, 1035.7, 777.0, 622.1.

**19. Synthesis of Ac-HHHHHHENLYFQGKGDPK(ac)K(ac)PRGKM-CO-SAL ester**

The Ac-HHHHHHENLYFQGKGDPK(ac)K(ac)PRGKM-CO-SAL was synthesized according to the **General Experimental Procedures 2.3.2.**

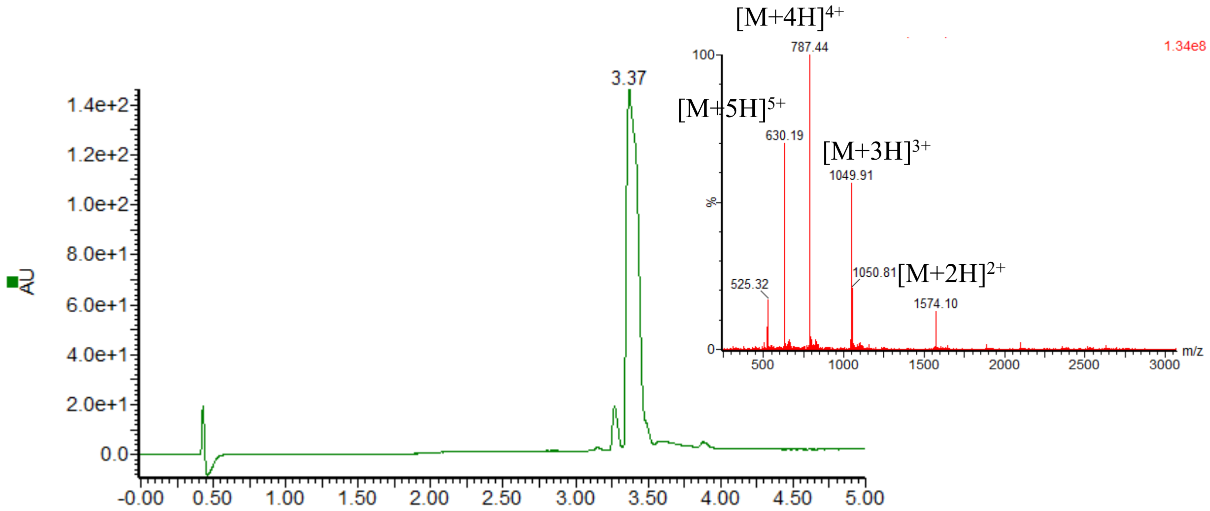


UV trace and corresponding MS from LC-MS analysis of purified product. ESI-MS calcd. molecular weight: 3146.51 $[M+2H]^{2+}$ m/z =1574.3, $[M+3H]^{3+}$ m/z = 1049.8, $[M+4H]^{4+}$ m/z = 787.6, $[M+5H]^{5+}$ m/z = 630.3, found 1574.1, 1049.9, 787.4, 630.2.

**20. Synthesis of Ac-HHHHHHENLYFQGKGDPK(ac)K(ac)PRGKM-CO-SAL ester**

The Ac-HHHHHHENLYFQGKGDPK(ac)KPRGK(ac)M-CO-SAL was synthesized according to the **General Experimental Procedures 2.3.2.**

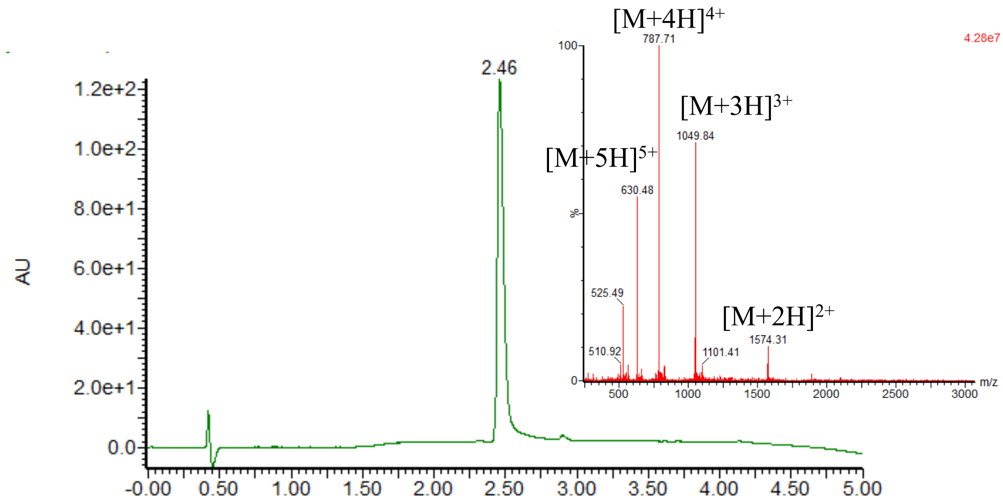


UV trace and corresponding MS from LC-MS analysis of purified product. ESI-MS calcd. molecular weight: 3146.51 $[M+2H]^{2+}$ m/z =1574.3, $[M+3H]^{3+}$ m/z = 1049.8, $[M+4H]^{4+}$ m/z =

787.6, $[M+5H]^{5+}$ m/z = 630.3, found 1574.3, 1049.9, 787.7, 630.5.

**21. Synthesis of Ac-HHHHHHENLYFQGK(ac)GDPK(ac)K(ac)PRGK(ac)M-CO-SAL ester**

The Ac-HHHHHHENLYFQGK(ac)GDPK(ac)K(ac)PRGK(ac)M-CO-SAL was synthesized according to the **General Experimental Procedures 2.3.2.**

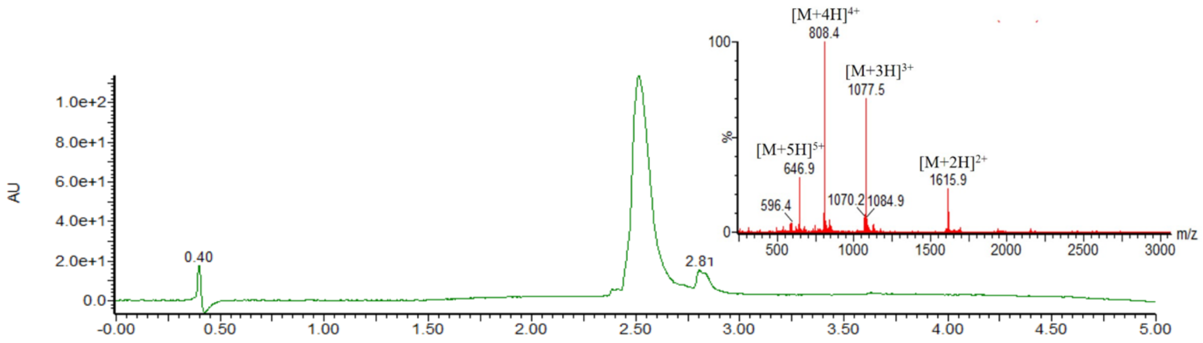


UV trace and corresponding MS from LC-MS analysis of purified product. ESI-MS calcd. molecular weight: 3230.43. $[M+2H]^{2+}$ m/z = 1616.2, $[M+3H]^{3+}$ m/z = 1077.8, $[M+4H]^{4+}$ m/z = 808.6, $[M+5H]^{5+}$ m/z = 647.1, found 1615.9, 1077.5, 808.4, 646.9.

**22. Synthesis of H-GKGDPKKPRGKMSS-OH**

The peptide was synthesized following standard SPPS protocol.

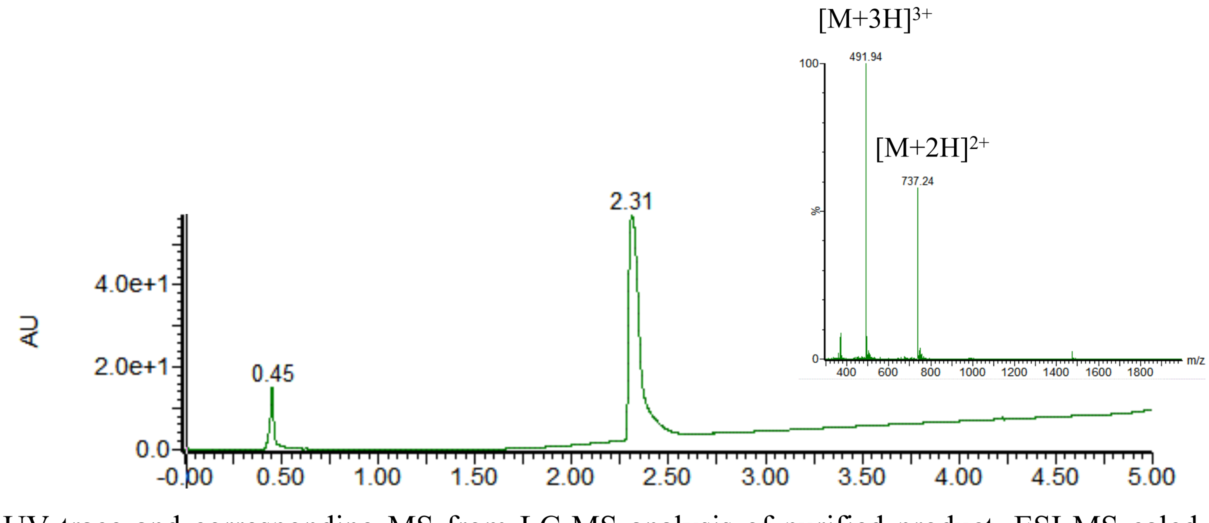


UV trace and corresponding MS from LC-MS analysis of purified product. ESI-MS calcd. molecular weight: 1472.73. $[M+2H]^{2+}$ m/z = 737.4, $[M+3H]^{3+}$ m/z = 491.9, found 737.2, 491.9.

**23. Synthesis of H-GK(ac)GDPKKPRGKMSS-OH**

The peptide was synthesized following standard SPPS protocol.

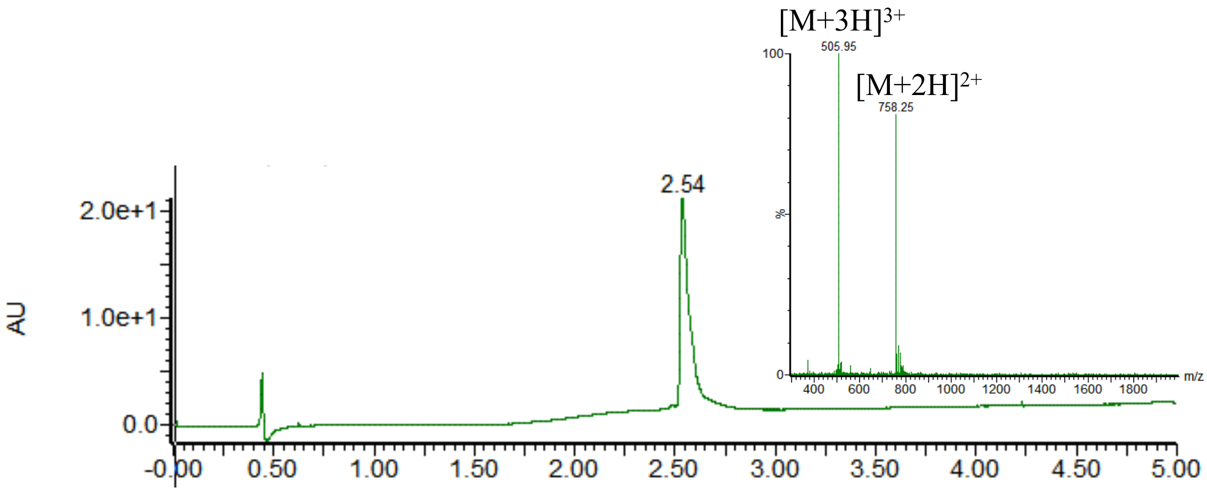


UV trace and corresponding MS from LC-MS analysis of purified product. ESI-MS calcd. molecular weight: 1514.77. $[M+2H]^{2+}$ m/z = 758.4, $[M+3H]^{3+}$ m/z = 505.9, found 758.3, 506.0.

##### 24. Synthesis of H-GKGDPK(ac)KPRGKMSS-OH

The peptide was synthesized following standard SPPS protocol.

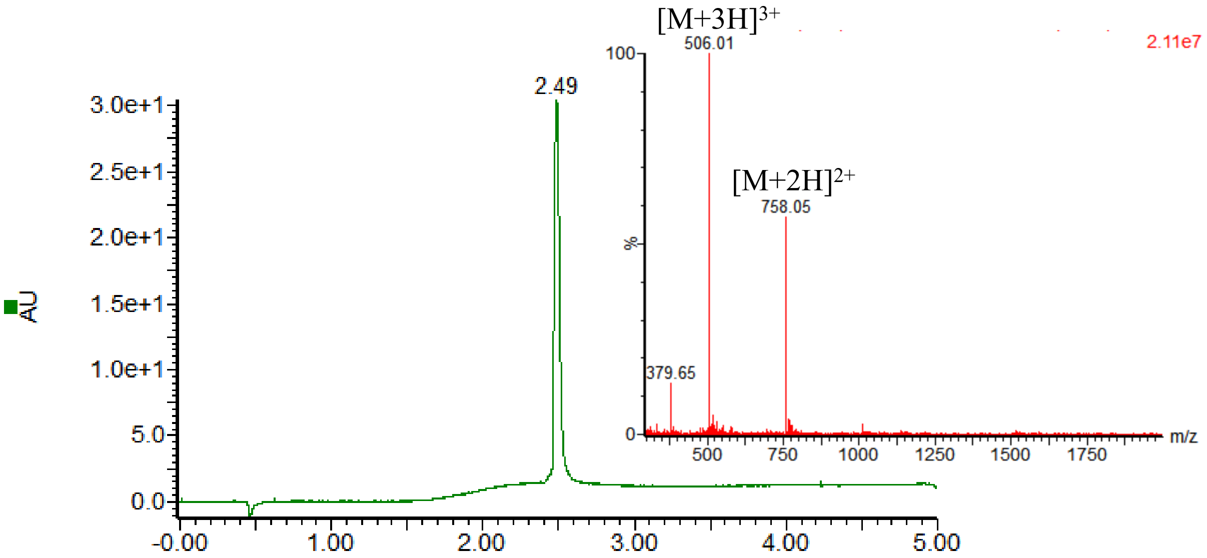


UV trace and corresponding MS from LC-MS analysis of purified product. ESI-MS calcd. molecular weight: 1514.77. $[M+2H]^{2+}$ m/z = 758.4, $[M+3H]^{3+}$ m/z = 505.9, found 758.1, 506.0.

##### 25. Synthesis of H-GKGDPKK(ac)PRGKMSS-OH

The peptide was synthesized following standard SPPS protocol.

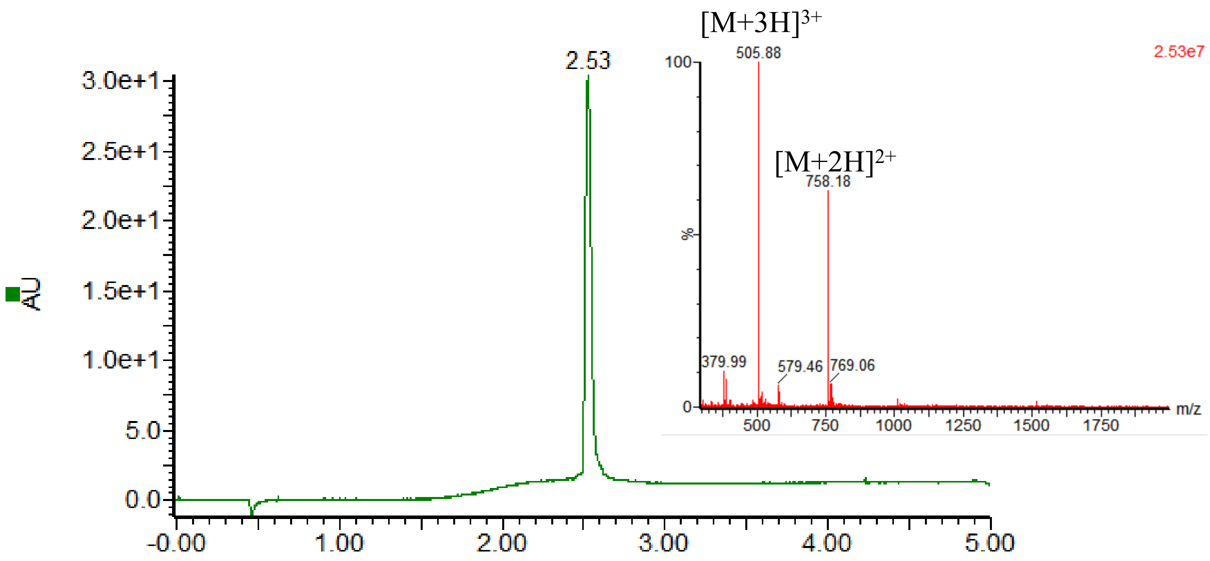


UV trace and corresponding MS from LC-MS analysis of purified product. ESI-MS calcd. molecular weight: 1514.77. $[M+2H]^{2+}$ m/z = 758.4, $[M+3H]^{3+}$ m/z = 505.9, found 758.2, 505.9.

**26. Synthesis of H-GKGDPKKPRGK(ac)MSS-OH**

The peptide was synthesized following standard SPPS protocol.

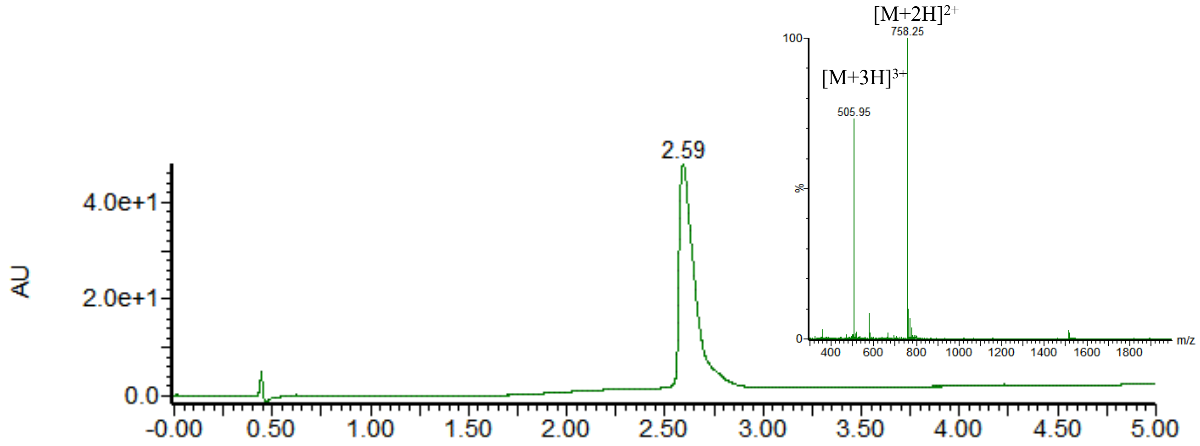


UV trace and corresponding MS from LC-MS analysis of purified product. ESI-MS calcd. molecular weight: 1514.77. $[M+2H]^{2+}$ m/z = 758.4, $[M+3H]^{3+}$ m/z = 505.9, found 758.3, 506.0.

**27. Synthesis of H-GKGDPK(ac)K(ac)PRGKMSS-OH**

The peptide was synthesized following standard SPPS protocol.

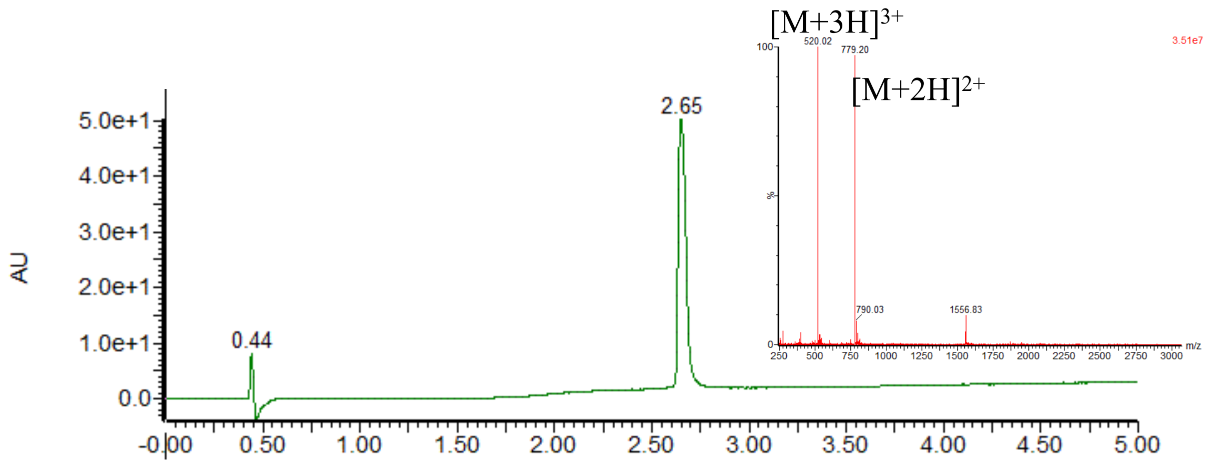


UV trace and corresponding MS from LC-MS analysis of purified product. ESI-MS calcd. molecular weight: 1556.81. $[M+2H]^{2+}$ m/z = 779.4, $[M+3H]^{3+}$ m/z = 519.9, found 779.2, 520.0.

**28. Synthesis of H-GKGDPK(ac)KPRGK(ac)MSS-OH**

The peptide was synthesized following standard SPPS protocol.

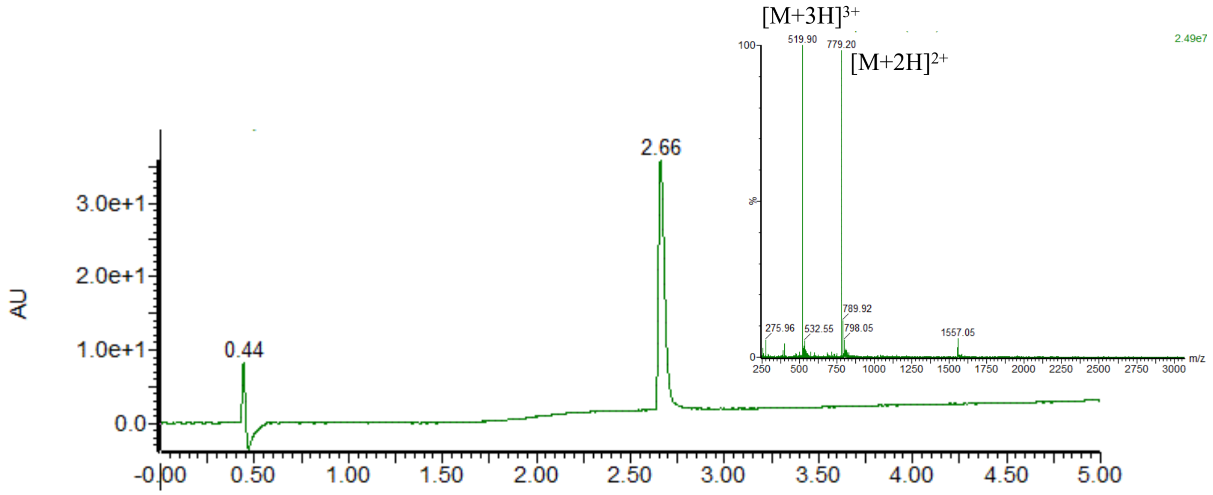


UV trace and corresponding MS from LC-MS analysis of purified product. ESI-MS calcd. molecular weight: 1556.81. $[M+2H]^{2+}$ m/z = 779.4, $[M+3H]^{3+}$ m/z = 519.9, found 779.2, 520.0.

**29. Synthesis of H-GK(ac)GDPK(ac)K(ac)PRGK(ac)MSS-OH**

The peptide was synthesized following standard SPPS protocol.

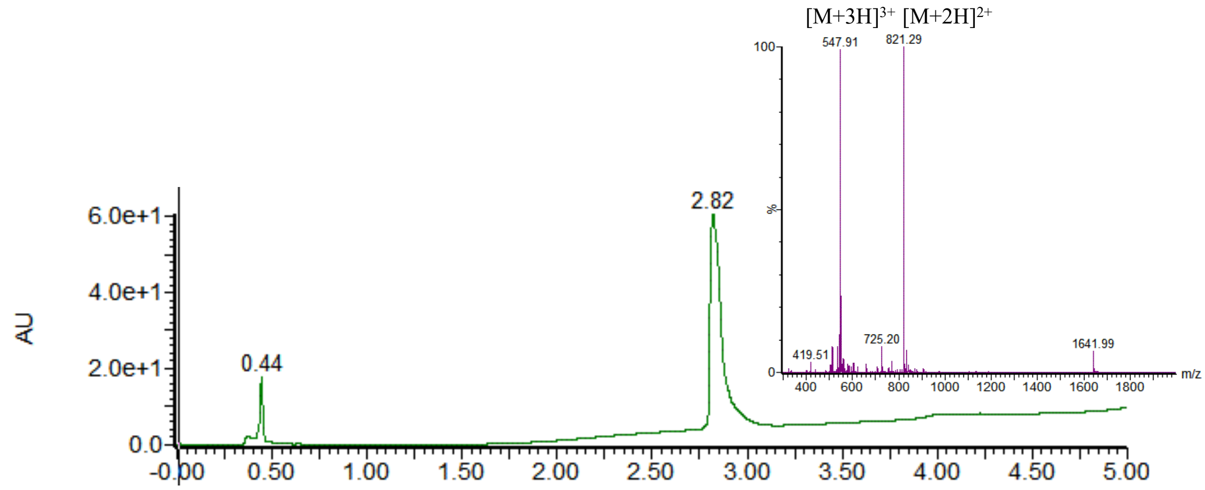


UV trace and corresponding MS from LC-MS analysis of purified product. ESI-MS calcd. molecular weight: 1640.88. $[M+2H]^{2+}$ m/z = 821.4, $[M+3H]^{3+}$ m/z = 548.0, found 821.3, 547.9.

**30. Synthesis of H-GKGDPQKPRGKMSS-OH**

The peptide was synthesized following standard SPPS protocol.

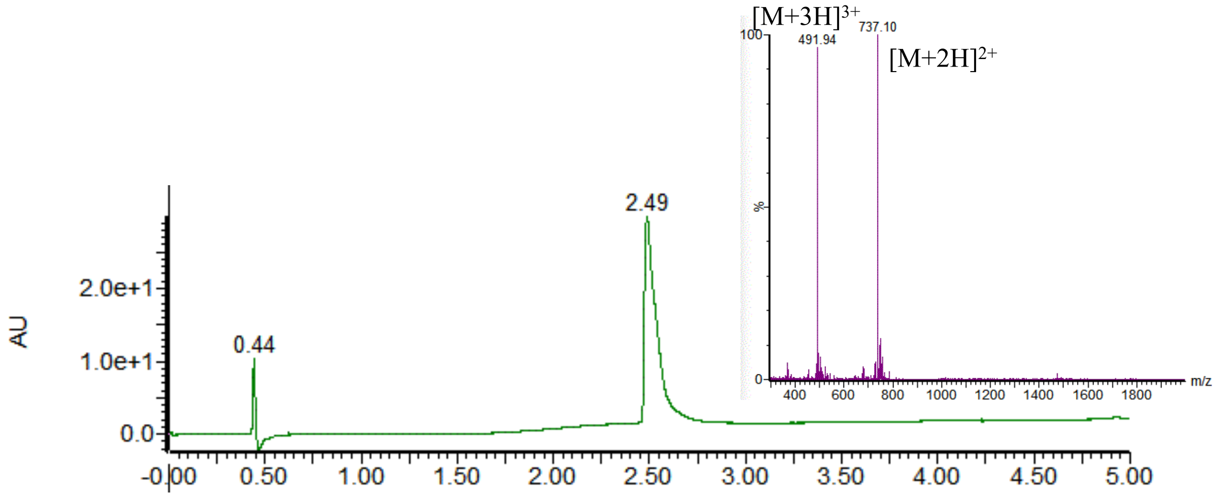


UV trace and corresponding MS from LC-MS analysis of purified product. ESI-MS calcd. molecular weight: 1472.69. $[M+2H]^{2+}$ m/z = 737.3, $[M+3H]^{3+}$ m/z = 491.9, found 737.1, 491.9.

**31. Synthesis of H-GKGDPKQPRGKMSS-OH**

The peptide was synthesized following standard SPPS protocol.

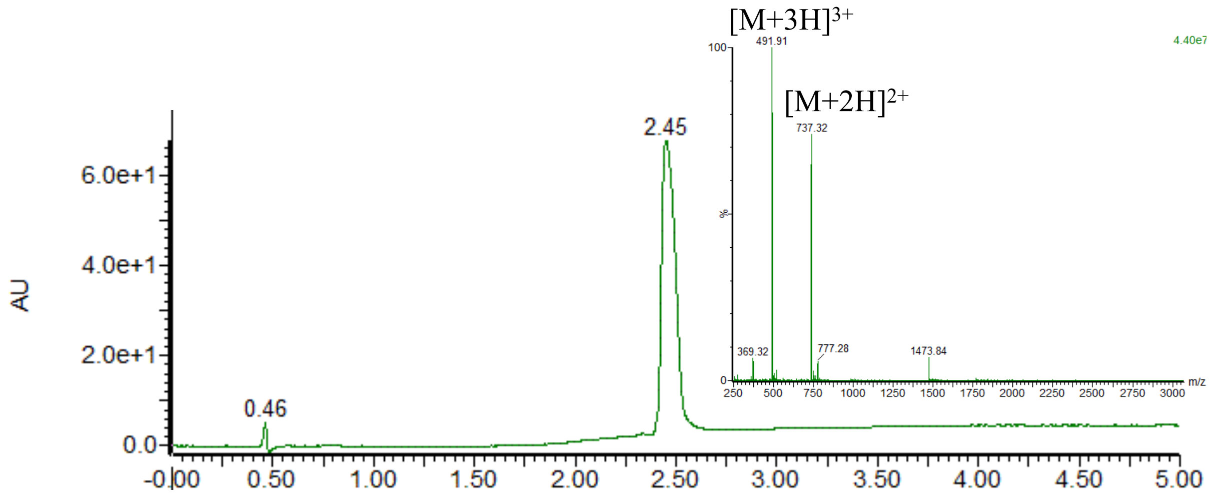


UV trace and corresponding MS from LC-MS analysis of purified product. ESI-MS calcd. molecular weight: 1472.69. $[M+2H]^{2+}$ m/z = 737.3, $[M+3H]^{3+}$ m/z = 491.9, found 737.3, 491.9.

**32. Synthesis of H-GKGDPKKPRGQMSS-OH**

The peptide was synthesized following standard SPPS protocol.

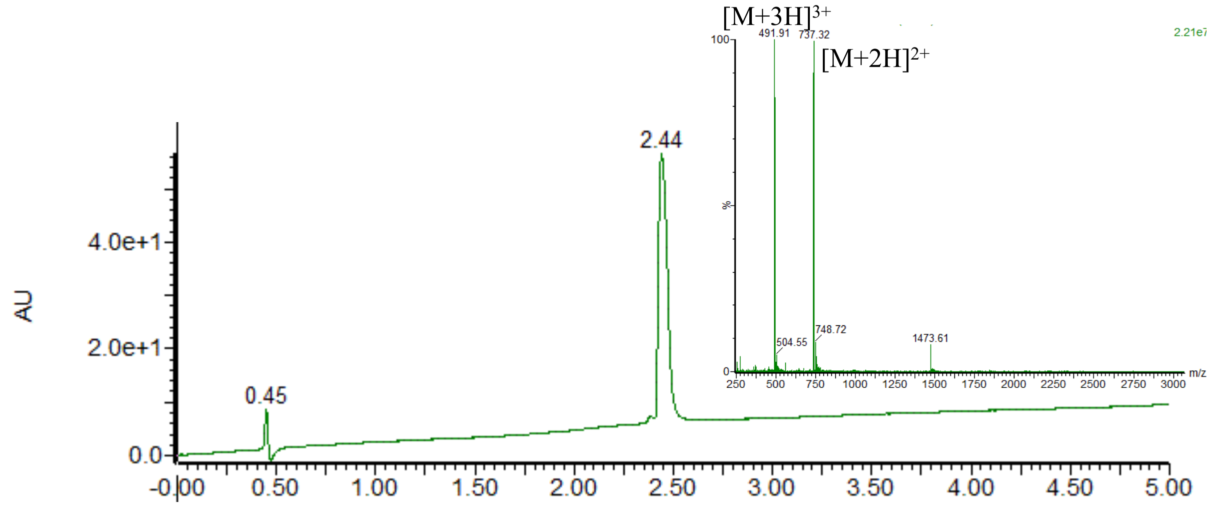


UV trace and corresponding MS from LC-MS analysis of purified product. ESI-MS calcd. molecular weight: 1472.69. $[M+2H]^{2+}$ m/z = 737.3, $[M+3H]^{3+}$ m/z = 491.9, found 737.3, 491.9.

**33. Synthesis of H-GKGDPQKPRGQMSS-OH**

The peptide was synthesized following standard SPPS protocol.

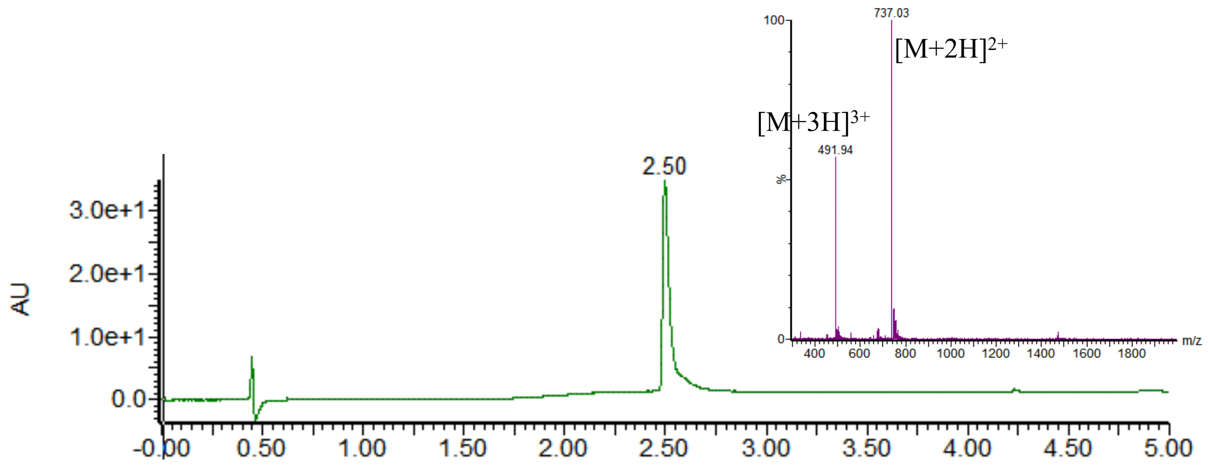


UV trace and corresponding MS from LC-MS analysis of purified product. ESI-MS calcd. molecular weight: 1472.64. $[M+2H]^{2+}$ m/z = 737.3, $[M+3H]^{3+}$ m/z = 491.9, found 737.0, 491.9.

**34. Synthesis of H-GKGDPKQPRGQMSS-OH**

The peptide was synthesized following standard SPPS protocol.

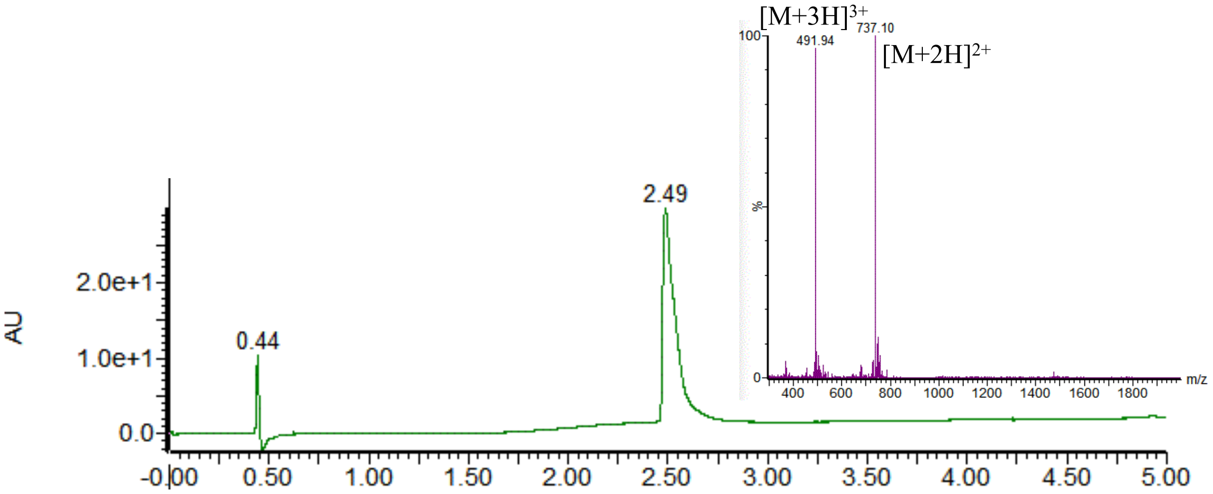


UV trace and corresponding MS from LC-MS analysis of purified product. ESI-MS calcd. molecular weight: 1472.64. $[M+2H]^{2+}$ m/z = 737.3, $[M+3H]^{3+}$ m/z = 491.9, found 737.1, 491.9.

2 Ito, T.; Kawahara, K.I.; Okamoto, K.; Yamada, S.; Yasuda, M.; Imaizumi, H.; Nawa, Y.; Meng, X.; Shrestha, B.; Hashiguchi, T.; Maruyama, I. Proteolytic cleavage of high mobility

group box 1 protein by thrombin-thrombomodulin complexes. *Arterioscler Thromb Vasc Biol.* **2008**, *28*, 1825-1830.

3 Gallwitz, M.; Enoksson, M.; Thorpe, M.; Hellman, L. The extended cleavage specificity of human thrombin. *PloS one* **2012**, *7*, e31756.
